# Localized calcium accumulations prime synapses for phagocyte removal in cortical neuroinflammation

**DOI:** 10.1101/758193

**Authors:** Mehrnoosh Jafari, Adrian-Minh Schumacher, Nicolas Snaidero, Tradite Neziraj, Emily M. Ullrich Gavilanes, Tanja Jürgens, Juan Daniel Flórez Weidinger, Stephanie S. Schmidt, Eduardo Beltrán, Nellwyn Hagan, Lisa Woodworth, Dimitry Ofengeim, Joseph Gans, Fred Wolf, Mario Kreutzfeldt, Ruben Portugues, Doron Merkler, Thomas Misgeld, Martin Kerschensteiner

## Abstract

Cortical pathology contributes to chronic cognitive impairment of patients suffering from the neuroinflammatory disease multiple sclerosis (MS). How such gray matter inflammation affects neuronal structure and function is not well understood. Here we use functional and structural *in vivo* imaging in a mouse model of cortical MS to demonstrate that bouts of cortical inflammation disrupt cortical circuit activity coincident with a widespread but transient loss of dendritic spines. Spines destined for removal show a local calcium accumulation and are subsequently removed by invading macrophages and activated microglia. Targeting phagocyte activation with a new antagonist of the colony-stimulating factor 1 receptor prevents cortical synapse loss. Overall, our study identifies synapse loss as a key pathological feature of inflammatory gray matter lesions that is amenable to immunomodulatory therapy.

**HIGHLIGHTS:**

- Widespread, but transient loss of synapses in inflammatory lesions and beyond
- Reversible impairment of neuronal firing and circuit function in the inflamed cortex
- Calcium dyshomeostasis of single spines precedes swift synapse loss
- Phagocyte-mediated spine pruning as targetable mechanism of synapse loss

## INTRODUCTION

Multiple sclerosis (MS) is a common and severe inflammatory disease of the central nervous system (CNS), clinically characterized by distinct phases. In most cases MS is initially dominated by bouts of acute neuroinflammation that result in formation of focal white matter lesions and can be targeted therapeutically (Reich et al., 2018). In contrast, the later progressive phase of the disease remains largely refractory to current therapeutic interventions and clinically appears reminiscent of classical neurodegenerative conditions. Indeed, it is increasingly recognized that pathology in the gray matter including the cortex is central to the progression of MS (Calabrese et al., 2015; Mahad et al., 2015; Ontaneda et al., 2017). Gray matter pathology correlates with accruing disability, predicts conversion to progressive MS and explains the increasing cognitive deficits that MS patients with advanced disease experience (Damjanovic et al., 2017; Eshaghi et al., 2018; Ontaneda et al., 2017; Scalfari et al., 2018). Despite this profound clinical importance, our understanding of the mechanisms underlying cortical pathology in progressive MS remains poor, in part because modelling gray matter inflammation has been difficult. Moreover, the neuroimaging and neuropathology approaches used thus far provide insufficient spatial and temporal resolution to unravel the neurodegenerative mechanisms resulting from cortical inflammation.

Gray matter pathology in progressive MS is characterized by inflammation dominated by innate immune cells, extensive loss of cortical myelin and some neuronal cell death (Lagumersindez-Denis et al., 2017; Mahad et al., 2015; Peterson et al., 2001). Of note, synapse loss might be a key neuronal pathology in such gray matter lesions (Dutta et al., 2011) as it occurs widespread throughout both the demyelinated, as well as the non-demyelinated, i.e. “normal-appearing” gray matter (Albert et al., 2017; Jürgens et al., 2016). Thus, in cortical MS, just as in classical neurodegenerative diseases, the synapse might be the initiation site of neuronal damage. However, thus far it remains unknown, whether such synapse loss in cortical MS has a local functional correlate, how it is mediated, whether it is reversible, and if or how it could be prevented by appropriate therapeutic intervention. *In vivo* imaging of the mouse cortex allows assessing local circuit function (Busche et al., 2019; Rose et al., 2016), the corresponding structural correlates (Xu et al., 2009) as well as the cellular mechanisms of neurodegeneration (Kuchibhotla et al., 2008; Merlini et al., 2019). With the recent development of rodent models of cortical neuroinflammation by others and us (Gardner et al., 2013; Lagumersindez-Denis et al., 2017; Lodygin et al., 2019; Merkler et al., 2006; Silva et al., 2018), imaging of MS-related pathology in cortical circuits has become possible. Here we interrogated the structural and functional neuronal consequences of such inflammatory changes using a range of *in vivo* imaging modalities. We show that widespread synapse loss is induced in the inflamed mouse cortex which mimics cortical MS lesions and surrounding gray matter. This synapse loss was present across cortical layers and accompanied by diminished neuronal activity. Once cortical inflammation receded, synaptic connectivity recovered and cortical neurons re-established their original firing patterns. Notably, in active inflammatory lesions single spines were primed for subsequent removal by lasting calcium overload – with the removal process being mediated by locally activated microglia, as well as infiltrating macrophages. Finally, blunting the activation, invasion and phagocytic activity of these immune cells by blocking signaling through the colony-stimulating factor 1 (CSF1) receptor protected synapses from removal, suggesting that phagocyte-mediated synapse pruning might be a target to prevent progression of cortical MS pathology.

## RESULTS

### Widespread synapse loss is induced in a mouse model of cortical MS

To induce neuroinflammation in the mouse cortex, we adapted previous targeted models of cortical MS pathology from rats (Gardner et al., 2013; Merkler et al., 2006). We stereotactically injected interferon gamma (IFNγ) and tumor necrosis factor alpha (TNFα), pro-inflammatory cytokines highly expressed in the meninges of progressive MS patients (Gardner et al., 2013; Magliozzi et al., 2018), into the somatosensory cortex of mice that were previously immunized with myelin-oligodendrocyte glycoprotein (MOG, **Figure 1A**). This resulted in the formation of cortical lesions that extended subpially to both hemispheres (**Figure 1B**) thus mimicking the widespread cortical pathology in progressive MS. Like their counterparts in humans (Lagumersindez-Denis et al., 2017; Lucchinetti et al., 2011), these cortical lesions were characterized by phagocyte activation and modest T cell infiltration causing extensive myelin loss with relative preservation of axons (**Figure S1**). To explore, whether such experimental gray matter lesions also reproduce the pervasive synaptic pathology observed in MS cortex (Jürgens et al., 2016), we reconstructed the dendrites of sparsely labeled layer III and layer V cortical projection neurons in *BiozziABH x Thy1-GFP-M* mice (**Figure 1C-1E**). We found that the density of dendritic spines was significantly reduced in both layers (**Figure 1F** and **1G**) at the acute stage of the cortical MS (c-MS) model (3 days post cytokine injection, 3dpci). Spine loss was observed ipsi- and contralaterally to the cytokine injection and was absent in mice that received either MOG immunization or cytokine injection alone (**Figure S2**). Notably, spine loss was reversible and spontaneously recovered over time (**Figure 1H**). Electron microscopy (EM) confirmed the transient reduction of cortical synapse density (**Figure 1I** and **1J**), indicating that synapse loss is not restricted to cortical projection neurons, but rather a global feature of the inflamed cortex.

**Fig. 1.**
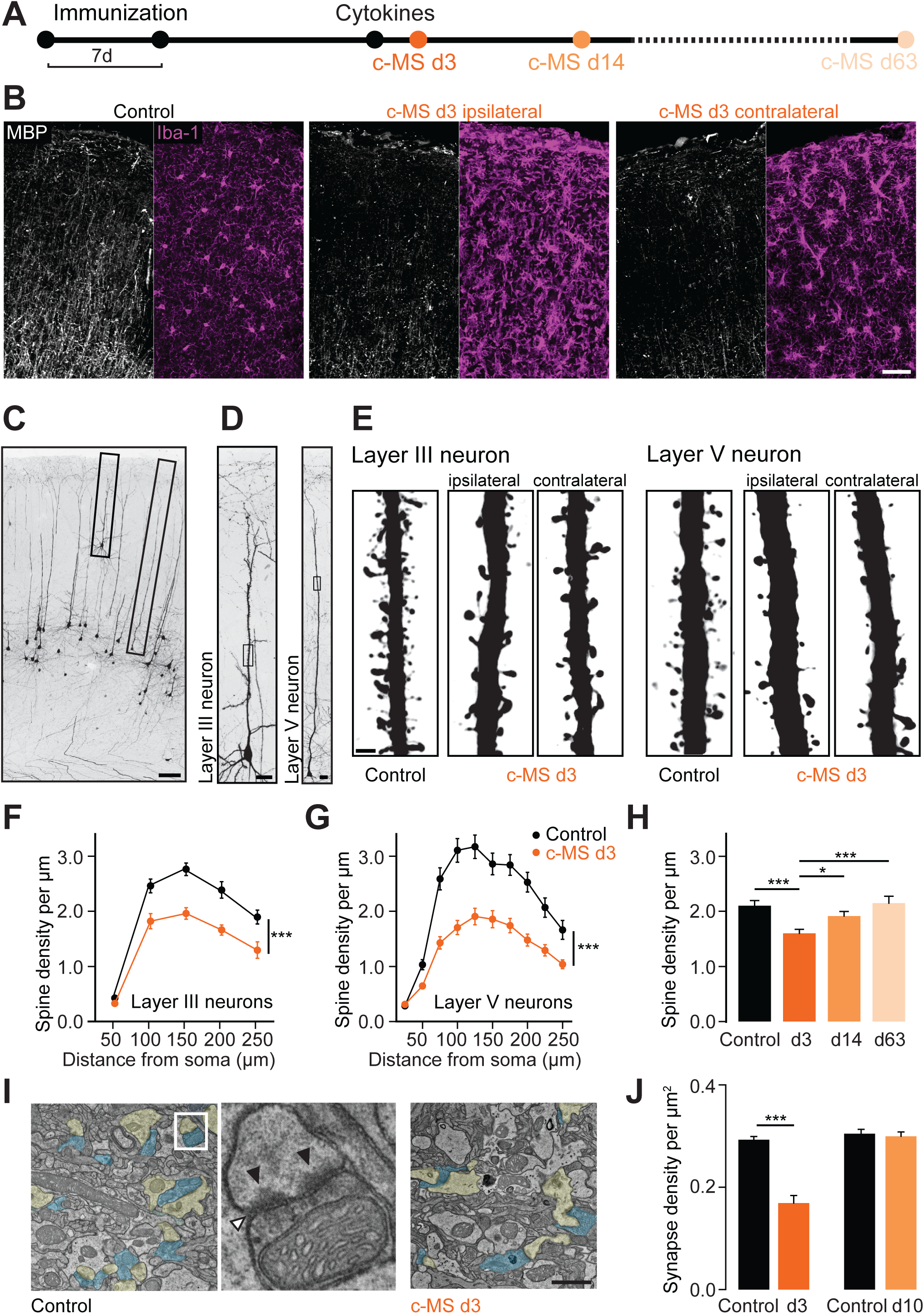
Projection neurons lose synaptic input in a model of cortical multiple sclerosis. **(A)** Schematic diagram of the experimental design to study spine density in the c-MS model at 3, 14 and 63 days after intracortical cytokine injection (c-MS d3, d14 and d63). **(B)** MBP staining (white) and Iba-1 staining (magenta) in EAE control (**left**) and c-MS model (**middle**, **right**). Demyelination and inflammation are observed both ipsilateral (**middle**) and contralateral (**right**) to the cytokine injection site in the c-MS model. **(C)** Confocal microscopy allows identification of pyramidal neurons located in cortical layers III and V in *BiozziABH x Thy1-GFP-M* mice. **(D)** Complete layer III and layer V neurons fully reconstructed using high-resolution confocal microscopy **(E)** Representative images of deconvoluted apical dendrites of layer III and layer V pyramidal neurons showing spines in healthy control and c-MS d3, ipsi- and contralateral to cytokine injection. **(F)** Spine density of layer III pyramidal neurons in healthy control and c-MS d3 along the apical dendrite (n=13 and n=15 neurons from n=6 mice per group, respectively; mean ± SEM). **(G)** Spine density of layer V pyramidal neurons in healthy control and c-MS d3 along the apical dendrite (n=21 and n=28 neurons from n=7 and n=8 mice, respectively; mean ± SEM). **(H)** Spine density of dendritic stretches of pyramidal layer V neurons in healthy control, c-MS d3, c-MS d14, and c-MS d63 (n=46, n=62, n=52 and n=24 dendritic stretches from n=7, n=8, n=8 and n=5 mice, respectively; mean ± SEM). **(I)** Electron micrographs of sham injected (**left**) and acute (d3) c-MS model (**right**) cortex layer 1, showing the synaptic compartments (presynaptic compartment in blue and postsynaptic in yellow). **Middle**: depiction of a synaptic complex with synaptic cleft (white arrowhead) and postsynaptic densities (black arrowheads). **(J)** Synapse density in cortical layer 1 in sham injected and c-MS animals obtained from electron micrograph analysis. Data shown as mean ± SEM (n=4 sham injected, n=4 c-MS d3 and n=5 sham injected, n=3 c-MS d10 mice). Scale bars in **B,** 50 µm; **C**, 100 µm; **D**, 20 µm; **E,** 2 µm **I**, 1 µm. Two-way RM ANOVA has been performed in **F** and **G**; One-way ANOVA followed by Bonferroni’s multiple comparisons test has been performed in **H** and **J**. *** P<0.001, *P<0.05.

### Cortical inflammation results in reversible neuronal silencing

To explore how this loss of synaptic input affects cortical function over time, we performed chronic calcium imaging in the somatosensory cortices of lightly anaesthetized mice using recombinant adeno-associated virus (rAAV)-expressed GCaMP6s (Chen et al., 2013). We recorded spontaneous calcium activity in populations of predominantly excitatory layer II/III neurons (96.6 ± 0.6%, GAD67 immunostaining negative, mean ± SEM, 1447 neurons from 4 mice analysed) tracked individually from before cortical lesion induction (4 days before cytokine injection), to the peak of synapse loss (3 dpci) and finally to the synapse recovery phase (10 and 17 dpci; **Figure 2A** and **Video S1**). We observed a shift towards lower frequencies of calcium transients with a significant drop in mean activity in acute gray matter lesions (**Figure 2B** and **2C**) consistent with neuronal silencing as a consequence of lost excitatory synaptic inputs (Montey and Quinlan, 2011; Rose et al., 2016). In line with this assumption, we observed a concomitant silencing and spine density reduction of the apical dendrites of layer V projection neurons that we recorded and reconstructed in *BiozziABH x Thy1-GCaMP6f* activity reporter mice (Dana et al., 2014) in the presence of cortical inflammation (**Figure S3**).

**Fig. 2.**
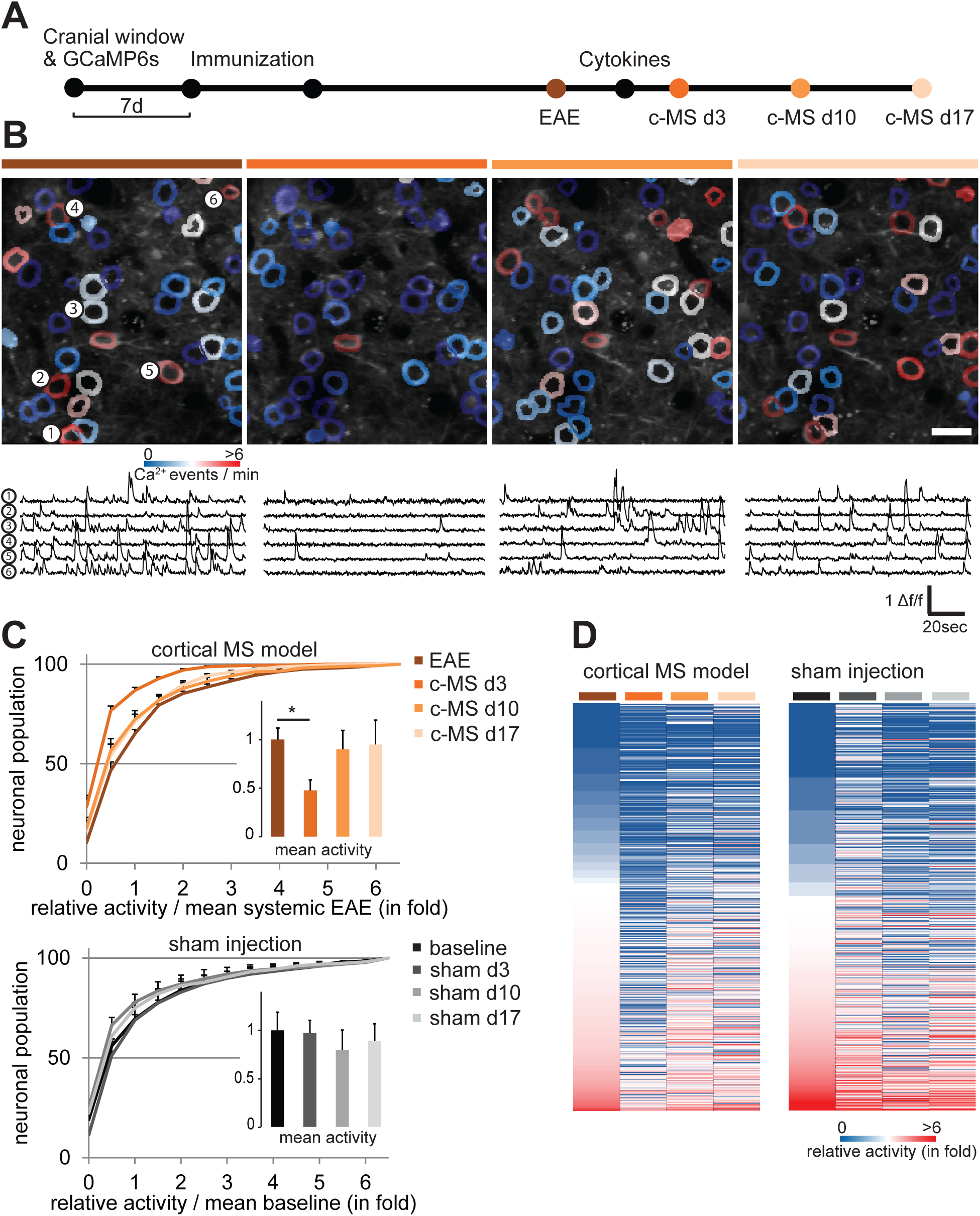
Neurons are silenced in a model of cortical multiple sclerosis. **(A)** Schematic diagram showing longitudinal *in vivo* imaging of neuronal activity in layer 2/3 somatosensory neurons in the c-MS model, using GCaMP6s encoded calcium indicator delivered via viral vector (AAV1.hSyn1.mRuby2.GCaMP6s). **(B)** *In vivo* multiphoton imaging of a layer 2/3 neuronal population over the course of cortical inflammation through a cranial window. **From left to right,** respectively: before cytokine injection (EAE), 3 days (d3), 10 days (d10) and 17 days (d17) post cytokine injection. **Top**: Grayscale images of GCaMP6s channel masked and color-coded for cytoplasmic Ca^2+^ events per minute. **Bottom**: Representative calcium activity traces displayed as delta f/f for the neurons marked in the left panel. **(C)** Cumulative plots of the neuronal activity for the entire population of neurons for each selected timepoint over the course of the c-MS model (**top**) and for sham injected animals (**bottom**). **Inserts**: Mean neuronal activity normalized to baseline for each timepoint shown as mean ± SEM (n=4 sham injected and n=5 c-MS mice, Shapiro-Wilk and one way ANOVA). **(D)** Heat map representation of the single neurons’ activities over the course of c-MS (**left**) and for the sham injected animals (**right**). Scale bar in **B**, 20µm. *P<0.01

Neuronal activity levels subsequently recovered in parallel with the restoration of synaptic connectivity (**Figure 2B** and **2C**). Activity changes are specific for the c-MS model, as no such alterations were observed in mice that received sham or cytokine injections only (**Figure 2B, 2C** and **Figure S4**). Notably, in contrast to activity and synapse loss, which recovered within two weeks, myelin loss only partially reverted over this time period as evaluated by spectral confocal reflectance microscopy (SCoRe) imaging (Schain et al., 2014) and ultrastructural analysis (**Figure S5**). Tracking activity levels of individual neurons over time further suggests that many neurons restore their pre-lesion activity level after transient silencing. For instance, neurons that were more active before lesion induction typically resumed a similarly high activity level after recovery (**Figure 2D**). To assess whether the recovery of individual firing patterns was due to a preset property of these neurons or the re-emergence of the pre-lesion micro-circuitry, we analyzed the structure of pairwise correlations of individual neuronal activity patterns over different imaging sessions before, during and after cortical neuroinflammation (**Figure 3A** and **3B**). Our results show that the session-to-initial session similarity (r_signal_) drops significantly during acute cortical inflammation (at 3 dpci) compared to sham-injected mice but then recovers during the resolution phase (at 10 and 17 dpci; **Figure 3C**). This indicates a remarkable resilience of neuronal circuit structure and activity despite transient inflammatory disruption in synaptic connectivity (Rose et al., 2016).

**Fig. 3.**
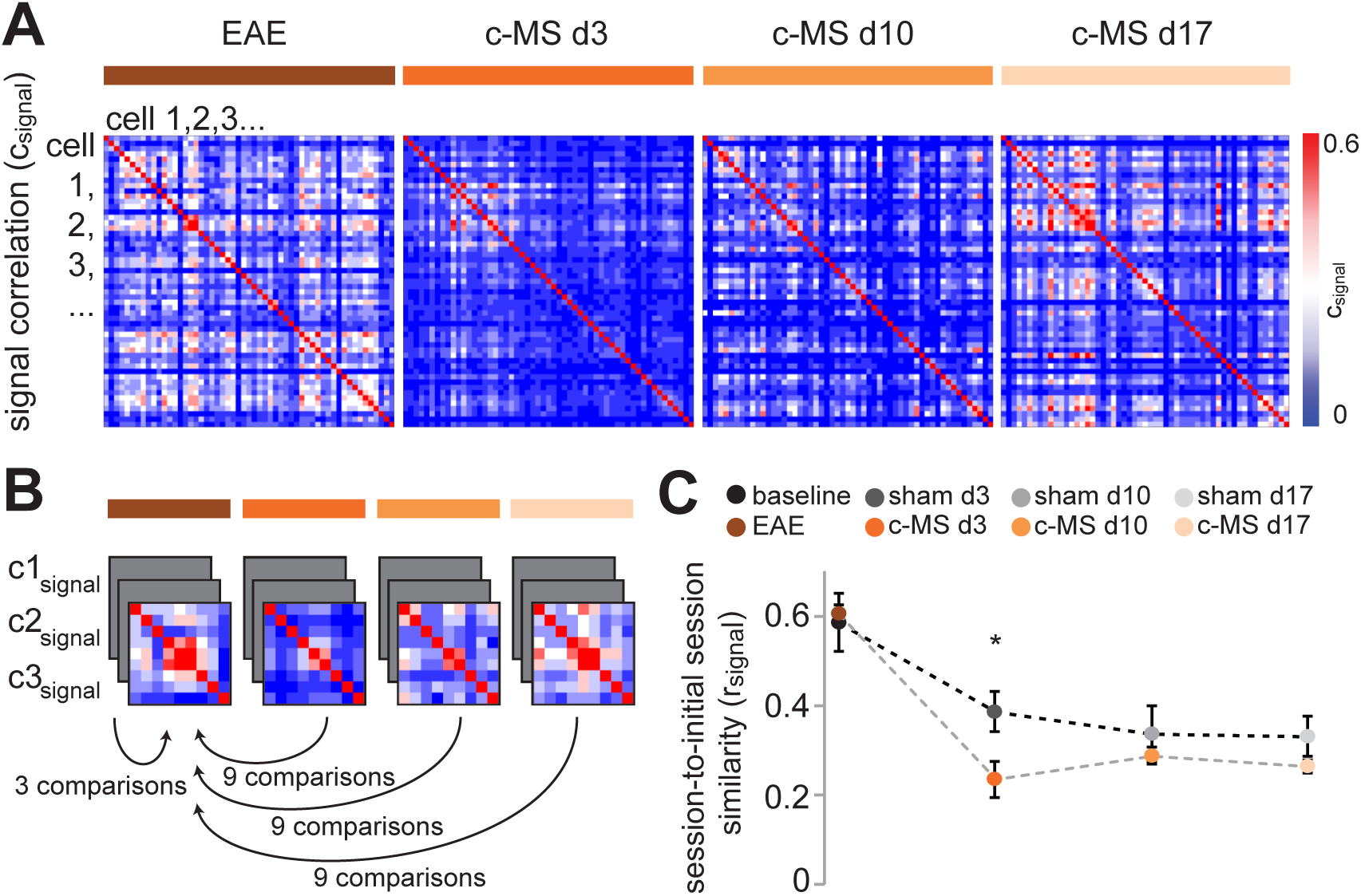
Resilience of neuronal circuit structure in a model of cortical multiple sclerosis. **(A)** Representative pairwise correlations of individual neuronal activity (c_signal_) in a somatosensory cortex area with 55 neurons over the course of the c-MS model in anaesthetized mice. **(B)** Schematic for session-wise similarity (r_signal_) calculation with c_signal_ structure at every timepoint (d3, d10 and d17) reported to before cytokine injection (EAE) in the c-MS model or to baseline in sham injected control. R_signal_ is calculated from the average of the multiple comparison based on three repetitive sessions at each timepoint over the course of the cortical MS model. **(C)** Average inter-timepoint similarity over the course of the c-MS model in comparison with sham injected animals. Data shown as mean ± SEM (n=4 control and n=4 c-MS, one-way ANOVA followed by t-test between c-MS and control). *P<0.05.

### Persistent local calcium accumulation predicts spine loss

Given its functional significance, we explored how synapse loss is induced in the inflamed gray matter and how such damage remains restricted to a subset of spines, while leaving the dendrite and many neighboring spines intact. Disturbances in calcium homeostasis have been shown to signal incipient loss of axons or dendrites (Kuchibhotla et al., 2008; Siffrin et al., 2010; Williams et al., 2014; Witte et al., 2019). Therefore, we explored whether local calcium dysregulation was confined to single dendritic spines by rAAV-mediated expression of the ratiometric calcium sensor Twitch-2b in a subset of cortical neurons (Thestrup et al., 2014). *In vivo* multiphoton imaging then allowed us to record spine calcium levels in the apical dendritic tufts at peak synapse loss (3 dpci) in the c-MS model (**Figure 4A** and **4B**). About 6 % of spines in the inflamed cortical tufts indeed displayed a localized calcium overload (> mean + 3 SD of control spines; **Figure 4C** and **4D**), while calcium levels in neuronal somata and the vast majority of dendrites remained unchanged (**Figure S6**). Notably, fast imaging of spine calcium dynamics revealed that high calcium spines showed a continuous calcium elevation that did not closely correlate to the local spine activity as high spike frequencies were observed both in calcium high and low spines (**Figure 4E**). These observations are consistent with a persistent local accumulation of calcium that occurs independently of altered synaptic activity.

**Fig. 4.**
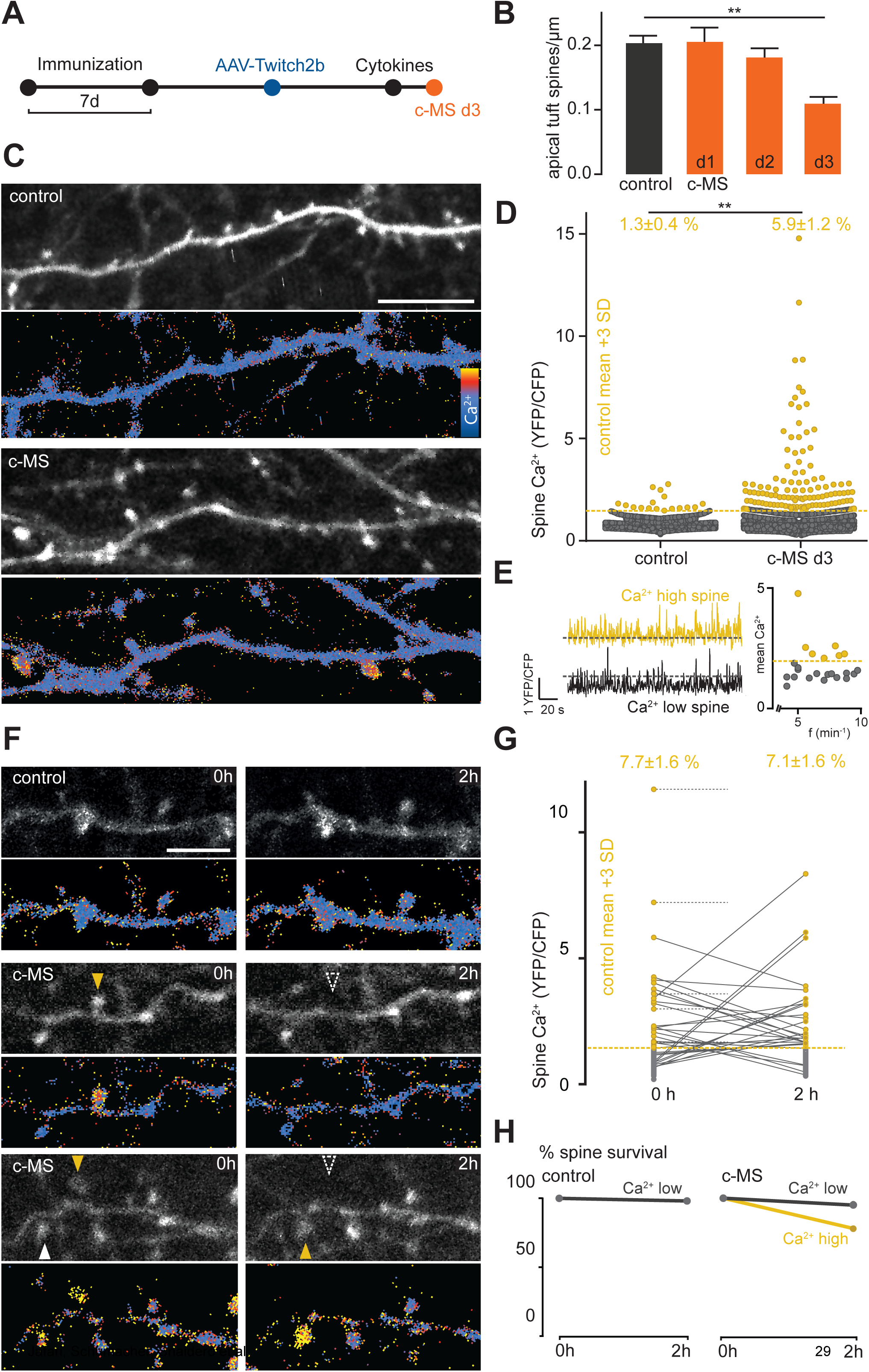
Localized calcium accumulations prime spines for removal. **(A)** Schematic diagram showing *in vivo* imaging of spine Ca^2+^ levels in the c-MS model using the FRET-based genetically encoded calcium indicator protein Twitch2b delivered via viral vector (AAV1.hSyn1.Twitch2b). **(B)** Timecourse of spine density in apical tuft dendrites of *BiozziABH x Thy1-GFP-M* mice, quantified *in situ* until 3d after induction of c-MS model (n=5 control, n=8 d1, n=7 d2 and n=7 d3, shown as mean ± SEM, 1way ANOVA). **(C)** *In vivo* multiphoton projection images of apical tuft dendrites with synaptic spines and their Ca^2+^ levels in healthy control (**top**) and acute (peak of disease, d3) cortical MS model (**bottom**). Grayscale images of YFP channel, ratiometric (YFP/CFP) images masked and color-coded for cytoplasmic Ca^2+^. **(D)** Ca^2+^ concentration of single spines in healthy and c-MS (d3) animals, plotted as YFP/CFP channel ratios (dashed line representing high Ca^2+^ threshold, 3SD above control mean) **Top:** Percentage of spines per animal with high Ca^2+^, shown as mean ± SEM (tested per animal in n=8 control and n=17 c-MS mice, Mann-Whitney U test). **(E) Left:** Representative calcium activity traces using resonant imaging, displayed as YFP/CFP normalized channel ratios over time in relation to high Ca^2+^ threshold (dashed lines). **Right:** mean baseline Ca^2+^ of single spines (normalized to control mean) shown in relation to their calcium activity (spiking frequency x min^−1^), dashed yellow line representing high Ca^2+^ threshold. **(F)** Multiphoton time-lapse images in healthy control and acute c-MS model (d2, d3) represented as a grayscale images of YFP channel and ratiometric (YFP/CFP) images masked and color-coded for cytoplasmic Ca^2+^. **Top:** stable control low Ca^2+^ spines. **Middle:** high Ca^2+^ spine (**left**, yellow arrow head) disappearing after 2h (**right,** dashed arrow head). **Bottom:** Spine (**left**, white arrow head) increasing volume and Ca^2+^ after 2h (**right**, yellow arrow head) next to neighboring spine with high Ca^2+^ disappearing after 2h (**right**, dashed arrow head). Gamma 1.2 for grayscale images **(G)** Ca^2+^ concentration of single spines in acute c-MS (d2, d3) animals over 2h, plotted as connected YFP/CFP channel ratios. Dashed lines represent high Ca^2+^ spines (yellow dots) disappearing within 2h. **Top:** Overall percentage of spines per animal with Ca^2+^ concentration > 3SD above control mean, shown as mean ± SEM (tested per animal in n=8 mice, paired t-test). **(H)** Percentage of surviving spines over time in healthy control animals (**left,** n=4 mice) and acute c-MS d2, d3 (**right,** n=8 mice). High Ca^2+^ spine cohort shown in yellow, neighboring Ca^2+^ spines in gray. Scale bars in **C**, 10 µm; **F**, 5 µm. **P < 0.01.

When we tracked the fate of these spines over time, about 25 % of the calcium-overloaded spines were lost within 2 hours, while on the same dendrite almost all spines that maintained calcium homeostasis persisted. During the same time period, however, several previously unaffected spines showed a rise in calcium that kept the fraction of calcium high spines stable (**Figure 4F–4H**). This suggests a highly localized and dynamic process of postsynaptic calcium overload that leads to the loss of ~1 % of cortical spines per hour. This rate can in little more than a day account for the total spine loss observed in the c-MS model (~30 %; **Figure 1** and **Figure 4B**). Thus, the local calcium overload of single spines appears to prime them for subsequent removal.

### Microglia and monocyte-derived macrophages execute synapse removal

We next assessed which cells execute this synapse removal. Mononuclear phagocytes are candidates, as the local activation of microglial and the infiltration of monocyte-derived macrophage coincides with synapse loss in the c-MS model (**Figure S7**). To evaluate the contribution of these cells to synapse loss, we quantified synapse phagocytosis by confocal 3D reconstruction of brain tissue from mice with genetically labeled phagocytes (microglia, CX3CR1^GFP^; invading macrophages, CCR2^RFP^) (Yamasaki et al., 2014) stained for pre-synaptic (Synapsin 1), post-synaptic (PSD95 and Homer) and lysosomal (LAMP 1) markers (**Figure 5A** and **5B**) and confirmed synapse uptake by EM (**Figure 5C**). Quantitative analysis showed a marked increase in engulfed pre- and post-synaptic material (both within and outside the lysosomal compartment) in microglial cells in the c-MS model compared to control gray matter (**Figure 5D, 5F** and **5G**). Furthermore, monocyte-derived macrophages contained pre and post-synaptic material in comparable amounts to microglial cells (**Figure 5E, 5H** and **5I**). Taken together these results indicate that local microglial cells and infiltrating monocyte-derived macrophages collaborate in the execution of synapse removal in inflamed gray matter.

**Fig. 5.**
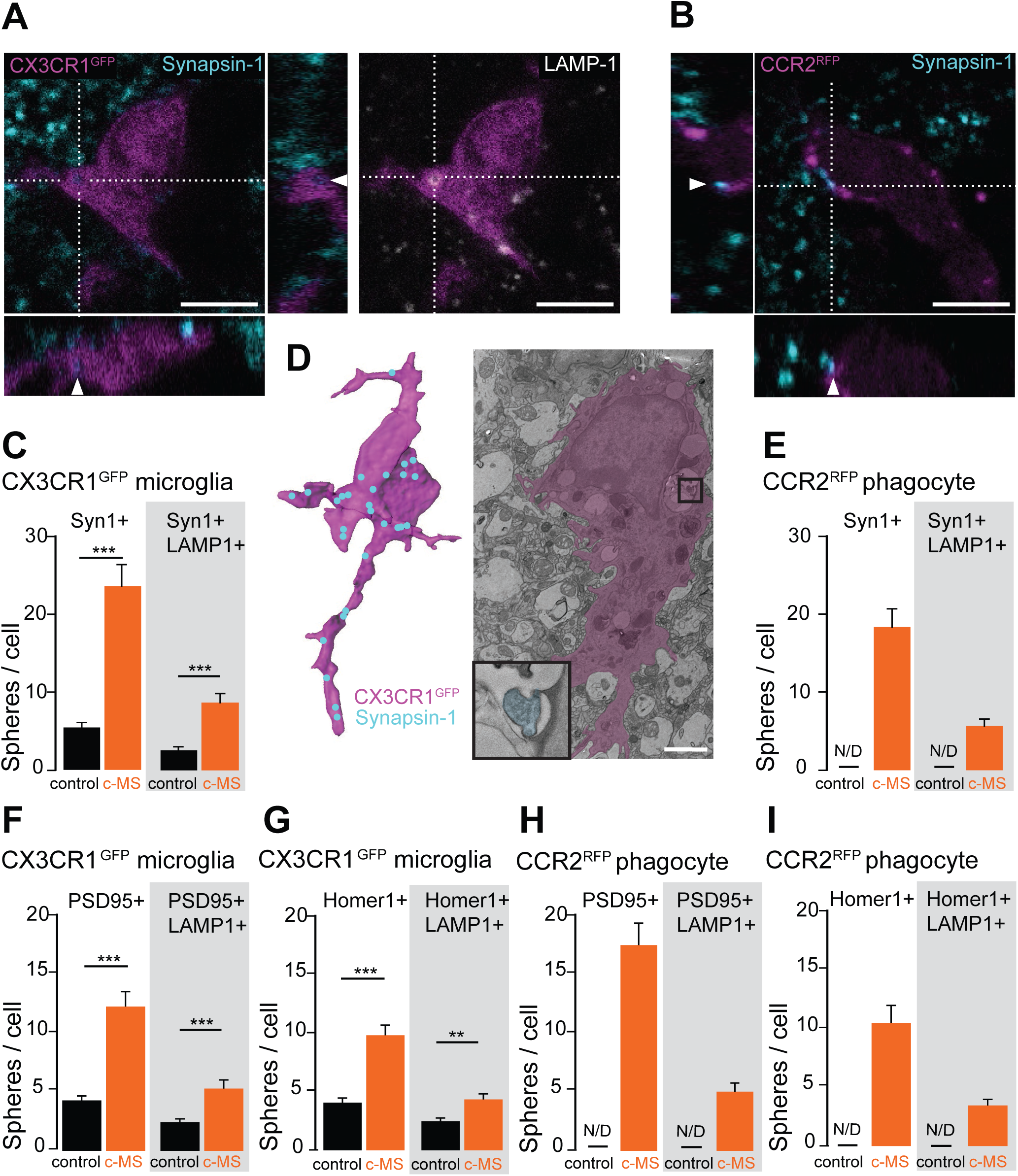
Mononuclear phagocytes remove synapses. **(A)** Orthogonal view of a CX3CR1^GFP^ cell (magenta) showing a Synapsin-1 inclusion (cyan, **left**) colocalized with LAMP-1 staining (white, **right**) suggesting removal of synapses by CNS resident microglial cells. **(B)** Orthogonal view of a CCR2^RFP^ cell (magenta) showing Synapsin-1 inclusion (cyan), suggesting removal of synapses by blood-derived mononuclear phagocytes. **(C)** Quantification of reconstructed Synapsin-1 positive spheres and Synapsin-1/LAMP1 double positive spheres inside CX3CR1^GFP^ cells in healthy control and c-MS d3 mice (n=40 cells from n=4 healthy control and n=5 c-MS d3 mice; mean ± SEM). **(D) Left:** 3D reconstruction of the cell in **A** (magenta) and its Synapsin-1 inclusions (cyan) with Imaris software allows studying synapse removal by mononuclear phagocytes. **Right:** Electron micrograph of a phagocytic cell (violet) in cortical layer 1 at the acute phase of c-MS. **Insert**: synaptic components (cyan) in a lysosomal compartment of the phagocyte cell. **(E)** Quantification of reconstructed Synapsin-1 positive spheres and Synapsin-1/LAMP1 double positive spheres inside CCR2^RFP^ cells in healthy control and c-MS d3 mice (n=0 cells from n=4 healthy control and n=50 cells from n=5 c-MS d3 mice; mean ± SEM). **(F, G)** Quantification of reconstructed spheres positive for postsynaptic markers (**F:** PSD95; **G:** Homer1) and double positive for LAMP1 inside CX3CR1^GFP^ cells in healthy control and c-MS d3 mice (n=50 cells from n=5 healthy control and n=5 c-MS d3 mice; mean ± SEM). **(H, I)** Quantification of reconstructed spheres positive for postsynaptic markers (**H:** PSD95; **I:** Homer1) and double positive for LAMP1 inside CCR2^RFP^ cells in healthy control and c-MS d3 mice (n=0 cells from n=4 healthy control and n=50 cells from n=5 c-MS d3 mice; mean ± SEM). Scale bars in **A** and **B**, 5 μm; **D**, 2 µm. Mann-Whitney U test in **C** and **E**; Unpaired two-tailed t-test in **F-I**. **P<0.01, *** P<0.001

### Targeted interference with CSF1 signaling prevents synapse loss in the c-MS model

Finally, we explored whether phagocyte-mediated synapse removal could be targeted therapeutically using a newly developed inhibitor of CSF1 signaling. While such inhibitors lead to the depletion of microglial cells at high concentrations (Elmore et al., 2014; Nissen et al., 2018), at lower concentrations, as used here they have been shown to inhibit phagocyte-mediated synapse removal in neurodegenerative disease models (Olmos-Alonso et al., 2016). Indeed, systemic application of the CSF1 receptor (CSF1R) antagonist during cortical lesion formation completely rescued synapse pathology in the c-MS model (**Figure 6A-C**). To further interrogate the mechanism underlying this effect, we FACS-sorted the following phagocyte populations from the inflamed CNS of *BiozziABH x CCR2^RFP/wt^* mice treated with either the CSF1R antagonist or vehicle solution (**Figure 6D, E**): CD45^int^ CD11b^int^ cells, which correspond to resting microglial cells, as well as CD45^high^, CD11b^high^ activated phagocytes. These we further differentiated in two subpopulations based on RFP expression, namely CCR2-RFP^positive^ cells, which likely represent recently infiltrated (monocyte-derived) phagocytes vs. CCR2-RFP^negative^ cells, presumed to be locally activated phagocytes (Saederup et al., 2010; Yamasaki et al., 2014). FACS analysis of these populations showed that CSF1R inhibition led to a selective reduction of the proportion of CCR2-RFP^positive^ CD45^high^ CD11b^high^ phagocytes isolated from the inflamed CNS, while the expression of activation markers was selectively reduced in the CCR2-RFP^negative^ CD45^high^ CD11b^high^ phagocytes. No obvious differences were observed in either the proportion or activation stage of resting microglial cells (**Figure S8**). RNAseq analysis of these cell populations further established that a marked change of the transcriptional profile in response to CSF1R inhibition was only apparent in CCR2-RFP^negative^ CD45^high^ CD11b^high^ phagocytes, while only few differentially regulated genes were detected in resting microglial cells (CD45^int^ CD11b^int^) or CCR2-RFP^positive^ CD45^high^ CD11b^high^ phagocytes (**Figure 6F-H**). Further analysis of the genes most-strongly regulated by CSF1R signaling in these locally activated phagocytes revealed that CSF1R blockade likely involves two collaborating mechanisms: First, a reduced expression of CCL2 (**Figure S8**), the major chemokine responsible for the infiltration of monocytes to the CNS (Huang et al., 2001). Indeed, a reduced number of monocyte-derived phagocytes was detected by histopathological analysis of cortical lesions derived from mice treated with the CSF1R antagonist, while numbers of microglial cells remained equal (**Figure 6K, N**), in line with the results of the FACS analysis (**Figure S8**). Second, CSF1R inhibition selectively altered the phenotype of the CCR2-RFP^negative^ CD45^high^ CD11b^high^ phagocyte population, as it diminishes APOE expression in these cells and thereby at least partially reverses the APOE-dependent disease-associated gene signature of these cells (Krasemann et al., 2017) towards a more homeostatic phenotype (**Figure 6I, J**). In line with such an altered phenotype, we also observed a reduced expression of scavenger receptors, such as the macrophage scavenger receptor 1 (MSR1) and the macrophage mannose receptor 1 (MRC1), as well as of genes such as CD68 and arginase-associated intracellular degradation pathways. To test whether this altered phagocyte phenotype would directly affect the capacity of local (but not infiltrating) phagocytes to remove synapses, we analyzed the lysosomal content and synapse uptake in *BiozziABH x CX3CR1^GFP/wt^* mice and *BiozziABH x CCR2^RFP/wt^* mice after CSF1R blockade. In line with the transcriptomic analysis, we found a selective reduction of lysosomal content and uptake of presynaptic material in CX3CR1-GFP expressing phagocytes that was not observed in the CCR2-RFP^positive^ cell population (**Figure 6L, M, O, P**). Taken together, this indicates that the primary cell population responding to the CSF1R inhibition are CCR2-RFP^negative^ CD45^high^ CD11b^high^ phagocytes. These mediate the therapeutic effect by reversing their disease-associated phenotype and ability to engulf synapses as well as dampening the recruitment of monocyte-derived macrophages (which are altered in number, but not in phenotype or synapse removal capacity). This dual mechanism of action thus controls both the central and the peripheral arm of the phagocyte response and effectively prevents synapse loss in the inflamed cortex.

**Fig. 6.**
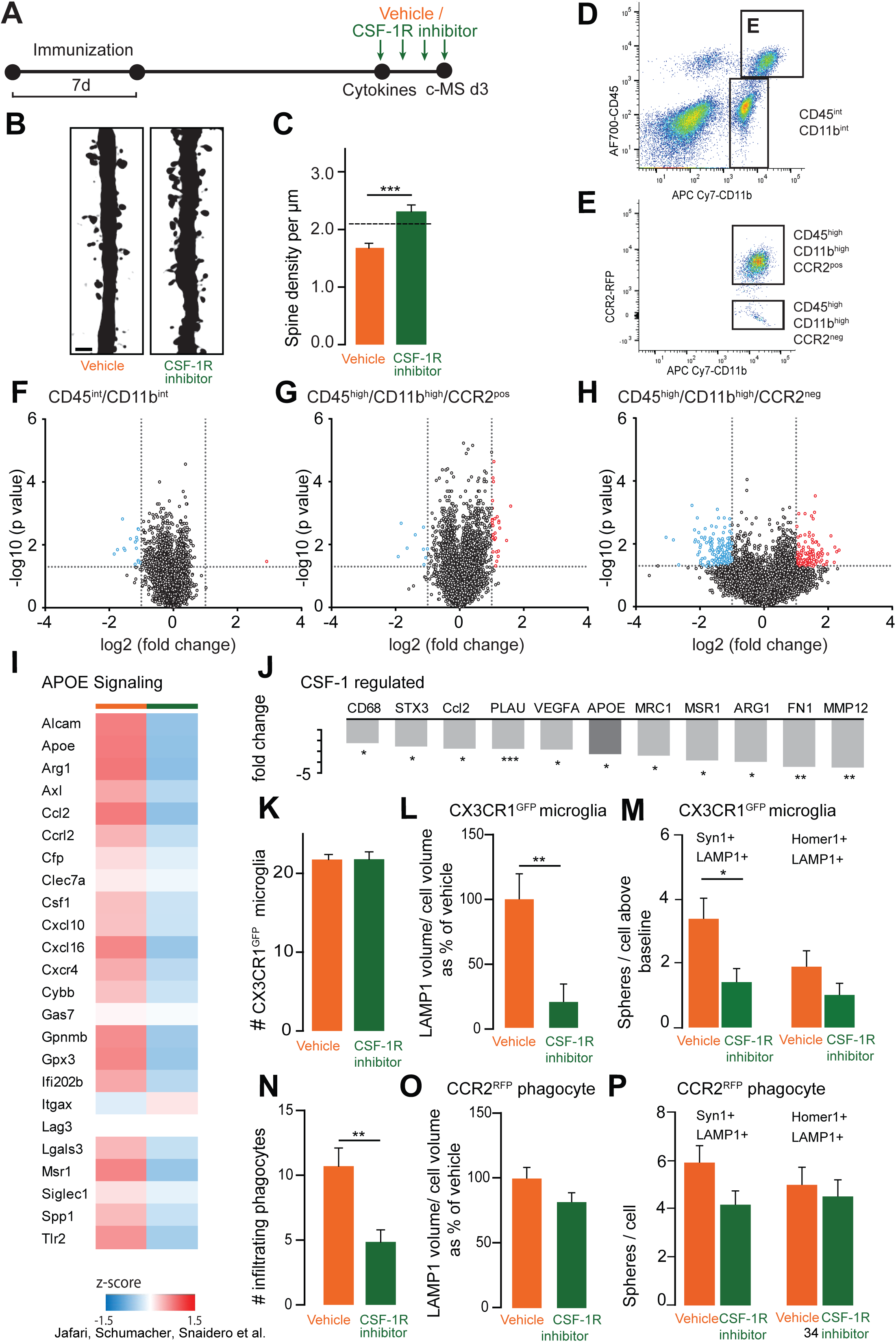
Blocking phagocyte entry and activation prevents synapse loss in the c-MS model. **(A)** Schematic diagram showing treatment with CSF-1R inhibitor or vehicle in the c-MS model. **(B)** Representative images of deconvoluted apical dendritic stretches from layer V pyramidal neurons showing spines in c-MS d3 mice treated with vehicle (**left**) or CSF-1R inhibitor (**right**) for 4 days. **(C)** Spine density of apical dendritic stretches from layer V pyramidal neurons in c-MS d3 mice treated with vehicle or CSF-1R inhibitor (n=65 and n=48 neurons from n=8 and n=7 mice; mean ± SEM). The dotted line shows the average level of spine density in healthy control mice based on Fig. 1G. **(D, E)** FACS analysis of cell populations in brains of c-MS model mice. **(D)** After separation of debris and doublets, live cells were sorted using CD45 and CD11b staining as gating strategies to identify mononuclear phagocytic cells. **(E)** CD45^high^ CD11b^high^ cells were further gated into two populations depending on the presence of CCR2-RFP positivity. **(F-H)** Volcano plot based on differentially expressed genes between CSF-1R inhibitor treatment vs. vehicle in CD45^int^ CD11b^int^ cells **(F)**, CD45^high^ CD11b^high^ CCR2^positive^ cells **(G)** and CD45^high^ CD11b^high^ CCR2^negative^ cells **(H)**. **(I)** Heatmap of treatment-affected genes associated with Apoe signaling as measured in transcriptomic analysis of CD45^high^ CD11b^high^ CCR2^negative^ cells (vehicle vs treatment, n=5 and n=6 mice respectively) **(J)** Genes predicted by Ingenuity Pathway Analysis to be directly regulated by CSF1 signaling (p-value cutoff of 0.05 and fold change cutoff of −2) in CD45^high^ CD11b^high^ CCR2^negative^ cells (vehicle vs treatment, n=5 and n=6 mice respectively) **(K)** Quantification of CX3CR1^GFP^ cells (CNS resident microglial cells) in cortical sections of c-MS d3 mice treated with vehicle or CSF-1R inhibitor (n=6 mice per group; mean ± SEM). **(L)** Normalized LAMP1 positive volume and **(M)** number of Synapsin-1/LAMP1 double positive spheres per CX3CR1^GFP^ cells in c-MS d3 mice treated with vehicle or CSF-1R inhibitor (n=5 mice per group; mean ± SEM). **(N)** Quantification of Iba-1^positive^ CX3CR1^GFP^ negative cells (infiltrating mononuclear cells) in c-MS d3 mice treated with vehicle or CSF-1R inhibitor (n=6 mice per group; mean ± SEM). **(O)** Normalized LAMP1 positive volume and **(P)** number of Synapsin-1/LAMP1 double positive spheres per CCR2^RFP^ cells in c-MS d3 mice treated with vehicle or CSF-1R inhibitor (vehicle vs treatment, n=4 vs n=5 respectively; mean ± SEM). Scale bar in **B**, 2 µm. Unpaired two-tailed t-test in **C, K-P.** *P<0.05 **P<0.01, *** P<0.001.

## DISCUSSION

While MS in its classical manifestation is dominated by white matter pathology and resulting sensory-motor symptoms, cortical symptoms, such as fatigue and subtle cognitive decline, also contribute to the patients’ loss of quality of life as the disease progresses (Calabrese et al., 2012; Rocca et al., 2014). The structural correlate of these symptoms is widespread gray matter pathology, with profound synapse loss, both in areas of gray matter demyelination (Dutta et al., 2011; Wegner et al., 2006) and beyond (Albert et al., 2017; Jürgens et al., 2016; Michailidou et al., 2015). Indeed, the degree of synapse loss in the MS cortex is comparable to that seen in early stages of classical neurodegenerative disorders, such as Alzheimer’s disease (DeKosky and Scheff, 1990; Terry et al., 1991). While much has been learned from applying *in vivo* imaging to disease models of such neurodegenerative conditions, both at the structural (Kuchibhotla et al., 2008; Merlini et al., 2019) and the functional level (Busche et al., 2008; Busche et al., 2019; Grienberger et al., 2012), our understanding of inflammatory gray matter pathology remains in its infancy. Thus, even very fundamental questions, such as whether structural and functional synapse changes are reversible, or which inflammatory mechanisms drive synapse loss remain largely unanswered.

The recent ability to model MS-related cortical neuroinflammation, using cortically targeted EAE models, now allowed us to address these questions. Indeed, our murine c-MS model recapitulates the widespread and pronounced cortical synapse loss also seen in MS (Albert et al., 2017; Dutta et al., 2011; Jürgens et al., 2016; Wegner et al., 2006). Importantly, this model also allows measuring the activity of local cortical circuitry by calcium imaging. Here, we observed hypoactivity of excitatory neurons (both in layer II/III and layer V) that tracks synapse loss of the same neuron populations. In contrast, in other settings of synapse loss, such as early amyloid pathology in a model of Alzheimer’s disease, hyperactivity has been described (Busche et al., 2008), which in part might be explained by substantial changes in overall cellular geometry due to dendrite remodeling (Siskova et al., 2014), which we found to be absent in cortical neuroinflammation. Furthermore, it is noteworthy, that in other models of Alzheimer’s disease-related pathology, which e.g. also incorporate tau aggregation, hypoactivity is also predominant (Marinkovic et al., 2019; Menkes-Caspi et al., 2015). In any case, prevailing evidence of functional imaging in human MS points to altered activity and circuit disconnection (Meijer et al., 2018; Rocca et al., 2010), relating to the findings at the cellular level in our model.

Given the profound structural and functional pathology that emerges from a single bout of cortical neuroinflammation, a remarkable observation of our study is the fact that such deficits can be entirely reversible once inflammation subsides. This speaks to a substantial endogenous neuronal repair potential in cortex, where apparently and in contrast to white matter also advanced structural alterations can still be reversed. Indeed once an axon has been transected in an inflammatory white matter lesion, no regeneration is possible, and compensatory remodeling is the prevailing, but limited mode of recovery (Kerschensteiner et al., 2004). In contrast, we now observe in the c-MS model that synapse loss, which includes removal of both pre- and postsynaptic material, is reversible without lasting structural imprint. Indeed, even the correlation landscape of local networks, likely indicative of regional interconnectivity or synchronized inputs, largely recovers, similar to what has been observed during reestablishment of visual cortex connectivity after sensory deprivation (Rose et al., 2016). Importantly, this degree of reversibility contrasts with the sustained damage apparent in human gray matter MS lesions. Beyond the obvious species difference, the two pathological processes differ in key respects - most importantly timing and local chronicity of an inflammatory environment. While the c-MS model involves only one episode of locally triggered neuroinflammation, driven by a single injection of cytokines, in MS sustained sources of such proinflammatory signaling exist, e.g. originating from lymphoid aggregates in the meninges, which correlate with increased levels of proinflammatory cytokines in the cerebrospinal fluid of corresponding patients (Magliozzi et al., 2018). However, such cytokines might well have complex effects, as for example TNFα can potentiate existing synaptic connections, and hence contribute to transient hyperactivity of cortical circuits, e.g. during the recovery phase of subcortical neuroinflammation (Ellwardt et al., 2018; Yang et al., 2013). Thus, other confounders, such as age (Confavreux and Vukusic, 2006; Scalfari et al., 2011) or threshold effects of accumulating subtle damage that exhaust repair or compensation capacities (Friese et al., 2014) might contribute to the lasting cortical deficits observed in progressive MS. Altogether, this argues that early intervention to prevent smoldering cortical inflammation from taking hold might allow endogenous recovery mechanisms to prevail and thereby counteract MS progression.

So what drives synapse removal and hence circuit dysfunction in cortical neuroinflammation? We find that phagocytes, including tissue-resident microglia and infiltrating (CCR2^positive^) macrophages engulf synaptic material. Such phagocyte-mediated synapse engulfment is also a hallmark of other neurodegenerative, neuroinfectious and neuropsychiatric disorders (Bialas et al., 2017; Di Liberto et al., 2018; Hong et al., 2016; Olmos-Alonso et al., 2016; Vasek et al., 2016), which stresses the unity of destructive mechanisms shared across a broad range of cortical pathologies. For driving cortical synapse removal, various mechanisms have been proposed. For instance, in development and neurodegeneration, it has been suggested that dysfunctional synapses are tagged by components of the complement system, which phagocytes recognize as ‘eat-me signals’ to initiate engulfment (Hong and Stevens, 2016). In other settings, such as a model of viral encephalitis, virally infected neurons express chemokines, such as CCL2 as ‘attack-me signals’, to attract the phagocytes that strip the neuron of its synapses (Di Liberto et al., 2018). To these executive mechanisms of phagocyte-mediated synapse stripping, we now add localized calcium dyshomeostasis as an early intrinsic feature of synapses that are fated for removal. We observe that individual spines transition into a lasting high-calcium state that predicts imminent removal. Such elevated calcium levels appear unrelated to synaptic hyperactivity (as we show by fast imaging), suggesting an intrinsic dysregulation of single spine calcium homeostasis, e.g. originating from local internal stores (Segal and Korkotian, 2014; Wu et al., 2017).

Importantly, irrespective of the precise mechanism of spine tagging, we identify CSF1 signaling as a possible therapeutic intervention point, which could remedy cortical synapse loss and is already under clinical investigation for neurodegenerative conditions (Olmos-Alonso et al., 2016). Notably, the dose of CSF1R antagonist treatment that we employ here avoids microglia depletion, which in chronic EAE models generates divergent outcomes (Nissen et al., 2018; Tanabe et al., 2019) and in a translational setting is hampered by the unpredictable risks of transiently ablating an essential CNS cell population. Non-depleting CSF1R-blockade, in contrast, appears to have a two-pronged beneficial effect in the context of our c-MS model - affecting the peripheral and the central arm of the phagocyte response in parallel. On the one hand, such CSF1R-blockade reduces infiltration of CCR2^positive^ phagocytes into the CNS, which is prominent in our model, but also in gray matter of MS patients (Lagumersindez-Denis et al., 2017). On the other hand, it reduces microglia-mediated synapse stripping inside the CNS, which is another feature shared by our model and gray matter pathology in progressive MS (Michailidou et al., 2015). The mechanism here, based on our transcriptomics analysis, is mediated primarily via locally activated phagocytes, which in response to CSF1R blockade alter their disease-associated phenotype, reduce the chemoattraction of peripheral phagocytes and down-regulate factors that affect endocytosis and engulfment suggesting a ‘paralysis’ of detrimental executive function. In summary, these data imply that also the progressive phase of MS, which appears refractory to current ‘peripheral’ immunomodulatory therapies (Hawker et al., 2009; Lublin et al., 2016; Wolinsky et al., 2007), might be amenable to a suitable strategy for CNS-targeted immunomodulation (Fox et al., 2018).

## ACKNOWLEDGEMENTS

We would like to thank A. Schmalz, M. Adrian, L. Schödel, B. Fiedler and Y. Hufnagel for excellent technical assistance, D. Matzek, B. Stahr, N. and M. Budak for animal husbandry. We acknowledge the Core Facility Flow Cytometry at the Biomedical Center, Ludwig-Maximilians-Universität München, for providing equipment, services and expertise.

Work in M.K.’s laboratory is financed through grants from the Deutsche Forschungsgemeinschaft (DFG; TRR128, Project B10 and B13), the European Research Council under the European Union’s Seventh Framework Program (FP/2007-2013; ERC Grant Agreement n. 310932), the German Federal Ministry of Research and Education (BMBF; Competence Network Multiple Sclerosis), the “Verein Therapieforschung für MS-Kranke e.V.”. Work in T.M.’s lab is supported by the DFG (CIPSM EXC114, CRC870, Mi 694/8-1), the German Center for Neurodegenerative Diseases (DZNE) and the European Research Council (FP/2007-2013; ERC Grant Agreement n. 616791). M.K. and T.M. are further supported by the DFG through a common grant (Ke 774/5-1/Mi 694/7-1) and the Munich Center for Systems Neurology (SyNergy EXC 2145). M.K. and M.J. are supported by a grant from the German National MS Society (DMSG). M.K., D.M. and T.M. were supported through grants from the Gemeinnützige Hertie Stiftung. Work in D.M.’s laboratory is further supported by the Swiss National Science Foundation (No. PP00P3_152928 and No. 310030_185321), the Klaus-Tschira Foundation, Helmut Horten Foundation and the Gebert-Rüf Foundation. Work in F.W.’s laboratory was financed through the DFG (CRC889, Project B6; CRC1286 Project C2), BMBF (01GQ0922), GIF (906-17.1/2006), VW foundation (ZN2632), BCCN (01GQ1005B) and the Max Planck Society. A-M.S. and T.N. were supported by the “*Förderprogramm für Forschung und Lehre*” at Ludwig-Maximilians University Munich.

## AUTHOR CONTRIBUTIONS

M.K., D.M. and T.M. conceived and designed the experiments. M.J., T.J., M.Kr. and D.M. established and characterized the c-MS model. M.J., A-M.S., E.M.U.G. and S.S. performed and evaluated cortical pathology in situ. A-M.S., T.N. and E.M.U.G. performed and evaluated *in vivo* calcium imaging of spines and dendrites. N.S., J.D.F.W. and F.W. performed and/or analysed neuronal activity patterns *in vivo*. N.S. performed electron microscopy analysis. R.P. and J.G. performed data analysis. M.J. and E.B. performed flow cytometry experiments. N.H., L.W. and D.O. developed and characterized the CSF1 receptor antagonist. M.J., A-M.S., N.S., D.M., T.M. and M.K. wrote the paper.

## STAR METHODS

### Lead Contact for Reagent and Resource Sharing

Requests for resources, reagents and further information should be directed to and will be fulfilled by Martin Kerschensteiner

### Experimental Model and Subject Details

#### Transgenic animals

All experiments were performed on animals with a F1 and F2 background of C57BL/6 and BiozziABH (strain designation BiozziABH/RijHsd, Harlan Laboratories), crossbred in our animal facilities. To assess dendritic and spine pathology in situ, we used *Thy1-GFP-M x BiozziABH* mice (derived from *Tg(Thy1-EGFP)MJrs/J*, gift from Joshua Sanes and Jeff Lichtman, Harvard University). To address phagocyte activation, infiltration and synaptic uptake, we used *CCR2^RFP^ x BiozziABH* animals (derived from *B6.129(Cg)-Ccr2^tm2.1Ifc^/J*, Jackson Laboratory) *and CX3CR1^GFP^ x BiozziABH* mice (*B6.129P-Cx3cr1^tm1Litt^/J*, Jackson Laboratory) or a F2 cross-breeding of both lines. For viral labelling, either *CCR2^RFP^ x BiozziABH* or *C57BL/6J x BiozziABH* were used. To assess calcium activity in layer 5 pyramidal neurons, we used *Thy1-GCamP6f x BiozziABH* mice (derived from *C57BL/6J-Tg(Thy1-GCaMP6f)GP5.5Dkim/J*, Jackson Laboratory, (Dana et al., 2014)). Mice were housed in individual ventilated cage systems under a 12:12 light/dark cycle, at a temperature of 22° ± 2°C and 55% ± 10% relative humidity with radiation-sterilized complete feed and sterilized water ad libitum. They were housed in social groups of a maximum of 5 mice in each standard housing cage and bedding. The cages were provided with enrichment consisting of play tunnels, nestlets to be used as nesting material and a red plastic mouse house. For husbandry, one male was housed with one or two females. Mice were weaned at postnatal day 21. Both female and male animals were included in experiments. All animal experiments were performed in accordance with regulations of the relevant animal welfare acts and protocols approved by the respective regulatory bodies.

### Method Details

#### Cortical multiple sclerosis model

Adult mice (6-14 weeks of age) were first induced with EAE as previously described. In short, we immunized animals with 200 μl of an emulsion containing 30 μg of purified recombinant myelin oligodendrocyte glycoprotein (MOG, N1-125, expressed in E. coli) and complete Freund’s adjuvant, consisting of incomplete Freund’s adjuvant (Sigma Aldrich) and 5 mg/ml mycobacterium tuberculosis H37 Ra (Difco). Pertussis toxin (100 ng, Sigma Aldrich) was administered intraperitoneally on day 0 and day 1 after immunization. The immunization procedure was repeated after 1 week. To evoke cortical lesions, we injected 2 µl of a cytokine mix of 0.25µg/µl TNF-alpha (R&D Systems) and 750 U/µl IFN-gamma (Peprotech) in PBS/0.1% BSA intracortically (coordinates: 1.2 mm lateral, 0.6 mm caudal to Bregma, depth 0.8 mm) 3 weeks after initial immunization. For all experimental animals, we assessed animal weight daily and assigned a score according to the severity of neurological deficits using an established EAE scoring scale: 0, no detectable clinical signs; 0.5, partial tail weakness; 1, tail paralysis; 1.5, gait instability or impaired righting ability; 2, hind limb paresis; 2.5, hind limb paresis with partial dragging; 3, hind limb paralysis; 3.5, hind limb paralysis and forelimb paresis; 4, hind limb and forelimb paralysis; 5, death.

#### Cranial window surgery

To gain optical access to the animal cortex to monitor neuronal activity over time and to assess *in vivo* calcium levels of dendrites and spines upon acute cortical inflammation, we performed a craniotomy and implanted a cranial window above the somatosensory cortex of the animals as previously described (Holtmaat et al., 2009). In brief, the mice were anaesthetized with a mixture of medetomidin (0.5 mg/kg), midazolam (5 mg/kg) and fentanyl (0.05 mg/kg) intraperitoneally. Craniotomy was performed using a 0.5 mm stainless steel drill head (Meisinger). For chronic imaging of neuronal activity, 0,5 µl of 10^12^ AAV1.hSyn.mRuby2.GSG.P2A.GCaMP6s (Vector Core, University of Pennsylvania) was injected 0.3 mm deep in the somatosensory cortex before sealing with a 4 mm cover class using dental cement. Mice were given buprenorphin (0.05 - 0.1 mg/kg) for analgesia every 12 h on the days following surgery. Under these conditions no increase in cellular toxicity due to viral burden was observed throughout the entire experiment timeline. Animals were first imaged 28 days after implantation of chronic cranial windows.

For acute *in vivo* imaging of baseline calcium levels of dendrites and spines, 0,5 µl of 10^12^ AAV1.hSyn1.Twitch2b.WPRE.SV40 (Vector Core, University of Pennsylvania) was injected 0.7 mm deep into the somatosensory cortex, 10d before the imaging timepoint. Analgesia was applied after surgery as described above. On the imaging day, acute cranial window implantation was performed as stated above.

#### Immunohistochemical stainings

For immunohistochemical analysis, mice were transcardially perfused with 4% (wt/vol) paraformaldehyde in phosphate buffered saline and after 12-24h post-fixation embedded in paraffin. Axon density was determined by Bielschowsky silver staining. Deparaffinized tissue sections were washed with distilled water and subsequently incubated in 20% silver nitrate solution for 20 min, followed by another washing step. Sections were incubated in 20% silver nitrate solution containing 5% ammonium hydroxide for 15 min in the dark. Sections were transferred in 0.6% ammonium hydroxide and swayed. 1/35 volume of the developer solution was added to the 20% silver nitrate solution containing 5% ammonium hydroxide while stirring. Subsequently the sections were transferred into the solution and developed until the color of the sections turned brown. Following washing in distilled water, sections were incubated in 2% sodium thiosulfate solution for 2 min.

For CD3 and Mac-3 stainings, we performed antigen retrieval of paraffin-embedded sections by treating rehydrated tissue sections for 30 s at 125 °C under pressure (20-22 psi) in 1x citrate buffer (pH = 6) in a Pascal pressure chamber. After cooling-down, antigen-retrieved sections were washed in distilled water and subsequently stained using a Biotin-Streptavidin System (Dako). All staining steps were performed at room temperature in a humid chamber. Sections were washed and blocked with peroxidase blocking solution (Dako) for 5 min. After two washing steps, sections were treated with the primary rat-antibody CD3 (1:100, Dako, A0452) and for Mac-3 (1:200, Biolegend, 108502). After washing, tissue sections were incubated with a biotinylated Rabbit Anti-Rat antibody diluted 1:250 in Real antibody diluent (Dako) for 15 min. After two washing steps, Real Streptavidin Peroxidase (HRP, Dako) was added for 15 min. Subsequently the sections were washed twice and processed with DAB-containing developer solution. Tissue sections were washed with water followed by counterstaining with Mayer’s Hämalaun.

MBP immunostainings were done using the DakoEnVision System at room temperature in a humid chamber. Sections were washed in 1x wash buffer and blocked with Real peroxidase blocking solution (Dako) for 5 min. After two washing steps, sections were treated with the primary rabbit polyclonal antibody (1:1000, Dako, A0623) diluted in Dako Real antibody diluent for 1 h. After washing, EnVision HRP-labeled polymer anti-rabbit IgG secondary antibody (Dako) was added for 30 min. Sections were washed twice and processed with DAB-containing developer solution (20 μl DAB chromogen + 1 ml DAB substrate buffer). Tissue sections were washed with water followed by counterstaining with Mayer’s Hämalaun. All stained sections described above were dehydrated and mounted in Ultrakitt for imaging.

#### Immunofluorescence stainings

Mice were transcardially perfused with 4% (wt/vol) paraformaldehyde in phosphate buffered saline. For dendritic reconstructions, myelin stainings and analysis of phagocyte activation and infiltration, brains were post-fixed for 12-24 hrs, isolated, and cut in 80-100 µm thick sections using a vibratome. Sections were first blocked with 10% goat serum (GS) in 0.5% (vol/vol) Triton X-100 in PBS (TPBS) and then incubated overnight with rabbit polyclonal MBP antibody (1:200, DAKO, A0623), rat anti-mouse I-A/I-E (1:300, BD Biosciences, 556999) or rabbit polyclonal Iba1 (1:300, Wako, 019-19741) in 1% GS / TPBS at 4° C. Stainings were visualized by incubating sections overnight with AlexaFluor-conjugated goat anti-rabbit (MBP) or goat anti-rat (MHC II) antibodies (1:500, Invitrogen) and counterstained with the Nissl-like nucleic acid stain NeuroTrace 435/455 (Invitrogen).

For evaluation of pre-synapse phagocytosis, brains were post-fixed 4-6 hrs, isolated, and cut in 40 µm thick sections using a vibratome. Sections were blocked as mentioned above and then incubated overnight with rabbit polyclonal Synapsin-1 antibody (1:500, Milipore, AB1543) and rat monoclonal LAMP-1 antibody (1:300, BioLegend, 121601) at room temperature. Stainings were visualized by incubating sections overnight with AlexaFluor-conjugated goat anti-rabbit (Synapsin-1) and goat anti-rat (LAMP1) antibodies (1:500, Invitrogen).

For evaluation of post-synapse phagocytosis, brains were fixed for 4-6 hrs, isolated and left in 30% sucrose for 3 days. The brains were then embedded in Tissue-Tek O.C.T. medium (Sakura Finetek Europe) and cut in 40 µm thick sections using a cryostat. Sections were blocked with 20% GS in PBS and incubated overnight with rabbit polyclonal PSD-95 antibody (1:200, Invitrogen, 51-6900) and rat monoclonal LAMP-1 antibody (1:300, BioLegend, 121601) or for 36 hours with guinea pig polyclonal Homer-1 antiserum (1:500, SySy, 160 004) and rat monoclonal LAMP-1 antibody (1:300, BioLegend, 121601) in 10% GS 0.3% TPBS. Stainings were visualized by incubating sections for 4 hours with AlexaFluor-conjugated goat-anti rabbit (PSD-95) or donkey anti-guinea pig (Homer-1) and goat anti-rat (LAMP1) antibodies (1:500, Invitrogen).

For the evaluation of excitatory and inhibitory neuronal fractions infected by AAV1.hSyn.mRuby2.GSG.P2A.GCaMP6s in the mice used in functional imaging, we perfused the animals as described above after the last imaging session. Brains were post-fixed for 12 hrs, isolated, and cut in 50 µm thick sections using a vibratome. Sections were first permeabilized in 2%Triton in PBS RT overnight then blocked in 2% FCS, 2% fishgelatine, 2% BSA in PBS for 2 hours at room temperature. Sections were then incubated overnight with mouse anti-GAD67 antibody (1:50, Millipore, MAB5406B), chicken anti-GFP (1:1000, Abcam, ab13970) in 10% blocking solution at 4° C overnight. Stainings were visualized by incubating sections for 2 hours with AlexaFluor-conjugated antibodies Alexa fluor 647 or Alexa fluor 488 (1:500, Invitrogen). All sections were mounted in Vectashield (Vector Laboratories).

#### Flow cytometry

For isolating the cells from blood, *CCR2^RFP^ x BiozziABH* mice were sacrificed and blood was collected from inferior vena cava and incubated with ACK lysis buffer (10:1, Gibco) for 5 minutes. For analysis of cell populations in the CNS, mice were transcardially perfused with ice-cold PBS directly after blood withdrawal and brains were isolated. Tissue was cut in small pieces and digested in RPMI containing 2% fetal calf serum (Sigma-Aldrich), 25mM HEPES (Sigma-Aldrich), DNase I (10ng/ml, StemCell Technologies) and Collagenase D (0,8mg/ml, Roche) for 30 min at 37°C. Digestion was stopped by adding 1:100 dilution of a 0.5M EDTA (Sigma-Aldrich) solution. After filtering the suspension through 70 µm cell strainers (Falcon), it was resuspended in a 30% solution of Percoll (Sigma-Aldrich). After 30 min of gradient centrifugation at 10.800 r.p.m., the myelin and red cells layers were removed and the remaining solution was filtered through 70 µm cell strainers (Falcon). Stainings were performed in ice-cold PBS after Fc-receptor blockade (CD16/32, 1:400, BD Pharmingen, 55314) using LIVE/DEAD staining (1:400, Invitrogen, L34957) and the following antibodies: Alexa Fluor 700 anti-mouse CD45 (1:400, Biolegend, 103128), APC-Cy7 rat anti-CD11b (1:300, BDPpharmingen, 557657), APC anti-mouse CD40 (1:100, BioLegend, 124612), PE/Cy7 anti-mouse CD86 (1:100, BioLegend, 105014) and BV711 rat anti-mouse I-A/I-E (MHC II, 1:200, BD Horizon, 563214). Samples were acquired on a FACSAria Fusion cell cytometer (BD Biosciences) and results analyzed by FlowJo software.

#### RNA Extraction, Library Preparation and Sequencing

For RNA extraction, samples were first randomized, then total RNA was extracted using the Picopure RNA Isolation kit (Thermo Fisher). RNA quantification and quality assessment was performed using a Nanodrop 8000 and RNA Pico 6000 (Bioanalyzer), respectively. Total RNA was diluted to 1 ng/ 10 µl of water prior to Oligo(dT) priming from Takara’s SmartSeq v4 Ultra Low Input RNA kit. Enriched mRNA was converted to full length cDNA then amplified using 11 PCR cycles (17 cycles for very low input samples). Final sequencing libraries were prepped and uniquely indexed using Illumina’s Nextera XT kit without modification to the standard protocol. Integrity of each sample library was assessed using the DNA High Sensitivity kit and run on a Bioanalyzer 2100. Library quantification was performed using Qubit then diluted to 1nM and pooled together. To prepare for sequencing, pooled libraries were denatured with NaOH then diluted to 1.8 pM according to Illumina’s denaturation protocol for a NextSeq500. Sequence runs were performed on an Illumina NextSeq500 using the High output kit and a 2×76bp paired end run.

#### Electron microscopy

For electron microscopy, perfusion was performed on anesthetized mice with 5 mL HBSS, followed by 30 mL fixative (2.5% glutaraldehyde, 4% paraformaldehyde in phosphate buffer). Brains were extracted and further post-fixed for 8 hours at 4°C in the same fixative. Afterwards, sagittal sections were done using a tissue slicer (Alto). The slice containing the cytokines (or sham) injection site were collected and further post-fixed with 2% OsO_4_ and 1.5% Ferocianide (Science Services), dehydrated by ethanol then acetone and Epon-embedded (Serva). 50 nm ultrathin sections from the area corresponding to the imaging site (2-3mm away from injection) were contrasted with 4% uranyl acetate (Science Services) and lead citrate (Sigma). The imaging was done on a TEM Jeol 1400+ (Jeol) equipped with a ruby camera (8M pixel, Jeol). The qualifications for the density of layer 1 myelin sheath was done within 60µm from the brain surface at 4000X on 720µm^2^ areas and the synaptic density was measured at the same depth at 8000X on 185µm^2^ areas.

#### Confocal microscopy

To image immunofluorescence stained tissue, we used upright confocal laser-scanning microscopes, FV1000 (Olympus) or SP8 (Leica), equipped with standard filter sets and laser lines. To reconstruct apical dendrites in *Thy1-GFP-M x BiozziABH* animals, 60x/1.35 NA oil immersion UPLSAPO Objective (Olympus) was used with a digital zoom of 3.5, Z-spacing of 200 nm and a pixel resolution of 75 nm pixel^−1^ to image 80-100 µm thick coronal brain sections. Images were deconvoluted using Huygens Essentials software. For the evaluation of synapse phagocytosis, 40 µm thick cryosections were imaged with a 40x/1.30 oil immersion HC PL APO CS2 objective (Leica), 3.0x zoom, Z-spacing of 200 nm and a pixel size of 97 nm pixel^−1^. To assess activation state and infiltration of phagocytes, 80 µm thick sections were imaged with a 20x/0.75 NA oil immersion HC PL APO CS2 objective (Leica) and 1.0x zoom in tile scan mode. Overview scans were overlayed on a scan of neurotrace signal, imaged with a 20x/0.85 NA oil immersion UPLSAPO Objective (Olympus).

#### *In vivo* imaging

For chronic *in vivo* imaging of neuronal activity, animals were first imaged 28 days after implantation of cranial windows. On the imaging day, animals were anaesthetized starting with 2% isoflurane and then placed on the imaging stage and provided with a constant flow of 1-1.5% isoflurane for the rest of the imaging period. The physiological state of the animals was continuously monitored with a MouseOx system (Starr Life Science Corp) equipped with a thigh sensor for small animals. Imaging of neuronal activity was performed on 2-3 areas of layer 2/3 somatosensory cortex at 15Hz with a resonant scanner (Olympus MPE-RS) using a femto-second pulsed Ti:Sapphire laser (Mai Tai Insight Deepsee, Spectra-Physics) with a maximum power of 35mW at the back focal plane. Each area was imaged 3 times 5 minutes within 1 hour. 25x/1.05 dipping cone water-immersion objective (Olympus) was used with a zoom of 1.5x and a pixel size of 660 nm pixel^−1^.

To image cortical myelin (layer 1) we performed spectral confocal reflectance microscopy (SCoRe) as described before. We used 3 fixed wavelength lasers (488, 539 and 633) to generate the SCoRe signal (adjustable bandpass filter: 488/4 and 539/4: Olympus, as well as a fix bandpass filter 636/8 BrightLine HC). Imaging was performed within the first 70 µm of the somatosensory cortex above the imaged areas used for the activity recordings. A 40x/0.8 water-immersion objective (Olympus) was used with a zoom of 1.0x and a pixel size of 200 nm pixel^−1^. Volume stacks penetrating 50-70 µm into the cortical layer 1 from the surface were acquired with a Z-spacing of 1 µm. A single plane from the stacks was used for each analysis.

For acute *in vivo* imaging of baseline calcium levels of dendrites and spines, animals were anaesthesized with a mixture of medetomidin (0.5 mg/kg), midazolam (5 mg/kg) and fentanyl (0.05 mg/kg) intraperitoneally and an acute cranial window was implanted as described above. After 30 min, *in vivo* microscopy was performed using a Olympus FV1200-MPE two-photon microscopy system, equipped with a femto-second pulsed Ti:Sapphire laser (Mai Tai HP-DS, Spectra-Physics) and laser power was attenuated by acousto-optical modulators. Emission was detected with non-descanned gallium arsenide phosphide (GaAsP) detectors. To image the genetically encoded calcium indicator Twitch2b, we used a two-photon wavelength of 840 nm to simultaneously excite both mCerulean3 and cpVenus^CD^. Fluorescence of mCerulean3 and cpVenus^CD^ was collected in a cyan channel (here referred to as “CFP”) and yellow channel (“YFP”), respectively, using emission barrier filter pairs with 455-490 and 526-557 nm. Images were acquired in 12 bit with an 25x/1.05 dipping cone water-immersion objective, pixel size of 124 nm pixel^−1^, dwell time of 2.0 μs pixel^−1^ and a laser power of 30-50 mW measured in the back focal plane. Volume stacks penetrating 30-60 µm into the cortical layer 1 from the surface were acquired with a Z-spacing of 1 µm.

Imaging of spine calcium activity was done using isoflurane anesthesia and a resonant scanner system at 15 Hz (Olympus MPE-RS) and continuous monitoring as stated above (section “chronic *in vivo* imaging of neuronal activity”). Images were acquired using a 25x/1.05 dipping cone water-immersion objective (Olympus) with a zoom of 4x and a pixel size of 249 nm pixel^−1^. Fluorescence of mCerulean3 and cpVenus^CD^ was collected in a cyan and yellow channel, respectively, using barrier filter pairs.

We observed no signs of photodamage in both healthy animals and c-MS animals over the imaging period using these imaging conditions. Animals showing signs of traumatic damage after window implantation were excluded from the analysis. We observed no significant difference in the fraction of spines with elevated calcium between the two stated methods of anaesthesia.

#### CSF1/LPS stimulation assay

Primary murine microglia were isolated with immunopanning and plated at 50k/well in PDL-coated 96-well plates. After resting overnight, cells were treated with CSF1R inhibitors (100nM, Sanofi) for 24 hours. CSF-1 (100ng/mL) or LPS (10ng/mL) was then added for an additional 24 hours and culture supernatants were collected and assayed for MCP-1 (CCL2) or IL-12p40 with ELISA.

#### Application of pharmacologicals

Cortical experimental autoimmune encephalomyelitis (c-MS model) was induced as mentioned above. Animals were treated daily from the day of surgery until perfusion (four days) with 20 mg/kg i.p. injection of vehicle or the novel CSF-1R inhibitor compound (Sanofi).

### Quantification and Statistical Analysis

#### Histopathological sections

From each staining, an image was chosen at the distance of 1700 to 2000 µm from the midline on both hemispheres using Pannoramic viewer software (3D-Histech). The sample images were rotated in Irfanview software and imported in FIJI software. The cortical layers I, II-IV and V were determined and cell counter plugin was used for evaluating each layer. For Bielschowsky’s staining, 2 parallel lines with a distance of 150 µm were located on each sample and the crossing filaments from each line was counted to evaluate axon density. For CD3 and Mac-3 staining, a ROI was drawn in each layer and the cells were quantified in each area. MBP areas were quantified applying a custom rule-set in Definiens Developer XD (Definiens AG, Version 2.7.0). Cortical regions were first manually selected and regions were rotated so that the cortical layers were horizontally oriented. Tissue was detected and cortical layers were automatically defined by their distance to the tissue-background border. Color deconvolution was applied to separate DAB and counterstain. MBP signals in the DAB channel were considered positive if above a defined intensity threshold. Total DAB+ area was reported for each region individually.

#### Confocal image processing and analysis

Post-processing of presented confocal images was done using the open-source image analysis software, ImageJ/Fiji (http://fiji.sc), and Photoshop (Adobe). Images were exported as RGB images to Photoshop and despeckled. Gamma corrections are stated in the figure legends, if done.

#### Reconstruction of cortical pyramidal neurons

Cortical pyramidal neurons were reconstructed using Volume Integration and Alignment System (VIAS) software (Rodriguez et al., 2003). Semi-automatic tracing of the apical dendrite and spines was performed by NeuronStudio software (Rodriguez et al., 2003; Wearne et al., 2005). For analyzing spine density over time and evaluating the therapeutic effects of CSF-1R inhibitor compound, dendritic stretches located at the layers III to IV with a dendritic radius of 0.55 to 1 µm coming from pyramidal layer V neurons were chosen. Neurons were located in the distance of 1000 to 2000 µm from the midline, contralateral to the injection site.

#### Analysis of neuronal activity

The analysis of neuronal activity recordings was performed with a custom software/graphical user interface. The software aligns the single frames of the data sets using a cumulative correlation maximization algorithm with the mRuby signal as reference. Using a calculated reference image (average frame, denoised running average, channel difference, coefficient of variation, between others) the software allows ROI definition manually or semi-automatically, providing an unique ID for each neuron. The ROIs defined at previous timepoints were imported into new recordings and automatically adjusted to compensate for the possible minor morphological changes. The pixels inside each ROI were averaged to extract a time signal of each neuron. The baseline signal F_0_ was calculated as the 10-percentile of the time series in a rolling window of 6 seconds. Signal was calculated using the following formula: dFF = (F-F_0_)/F_0_, which if necessary was corrected for neuropil contamination by subtracting a fraction (usually 60%) of the dFF calculated using a background ROI surrounding the neuron and excluding any other marked cell. The final dFF was processed with an exponential filter with a time constant of (0,01). Calcium events for the recording of each neuron were calculated using a non-negative deconvolution algorithm (Vogelstein et al., 2010), where the resulting likelihood per frame is processed to separate the individual events by summation and thresholding. The final neuronal activity for each neuron was calculated from the average of 3 recordings (see *in vivo* imaging section).

#### Neuronal population correlation analysis

At each timepoint, every region was imaged for three sessions (5 minutes/session), and a signal correlation matrix was obtained for each session. The correlation similarity between any two such matrices is defined as the correlation of the individual entries in the upper-triangular part of the correlation matrices (Rose et al., 2016). To compare the population dynamics at any given timepoint to the initial population dynamics, the similarity between all correlation matrices at the given timepoint and all correlation matrices in the initial session were calculated and the resulting values averaged. For the c-MS d3, c-MS d10 and c-MS d17 timepoints this involves averaging nine similarities (the three signal correlation matrices at each timepoint compared to the three initial signal correlation matrices). For the initial session this similarity involved three comparisons.

#### Dendritic and spine calcium levels

For analysis of Ca^2+^ levels in spines and dendrites using the Twitch2b sensor signal, the CFP and YFP (FRET) channels were individually visualized in a greyscale look-up table to determine the region of interest. Fluorescence intensities of the spine head or 5 regions in the dendrites shaft were measured in the CFP and YFP channels, respectively, and nearby non-neurite areas were selected for background correction. As we detected crosstalk from the CFP to the YFP channel (not vice-versa), we corrected the YFP signal by subtracting the measured crosstalk-fraction of the CFP signal. Background-corrected YFP/CFP ratios were interpreted as a proxy of Ca^2+^ concentration as previously established for the used FRET-based calcium sensor. We excluded spines that exhibited a signal-to-noise ratio < 5.2, as measured in YFP channel. Spines or dendrites were considered to be Ca^2+^-elevated if YFP/CFP ratios showed elevation greater than mean plus 3 standard deviations of the ratios measured in healthy control animals. Population analysis of spine and dendritic Ca^2+^ was done by a blinded investigator. For dynamic analysis of spine fate, per region 10 dendrites including all spines were identified in both timepoints and measured.

For the assessment of spine calcium activity, high-frequency resonant images were acquired (see *in vivo* imaging section) and registered using a custom MATLAB script. Spine and nearby non-neurite background ROIs were manually defined and measured over time in the CFP and YFP channels as stated above. Background-corrected YFP/CFP ratios were interpreted as a proxy of Ca^2+^ concentration. Raw values were smoothed by removing single outlier values (negative, >15) and using a 3-neighboring average algorithm. Baseline drift was assessed using a 10s moving average and, when no drift was observed, overall baseline mean and standard deviation calculated. Spiking activity was defined as above-threshold activity (baseline mean plus 2 SD) for > 0.25s and calculated as frequency (min^−1^).

Ratiometric images presented in this work were processed as follows: in Fiji, intensity projections of 3D stack sections were created from CFP and YFP images individually and a binary thresholded mask of dendritic outlines was generated from the image channel with higher signal-to-noise (after application of pixel outlier filters). Projection images were each multiplied by the binary mask and the resulting images divided by each other (YFP/CFP). The resulting image was pseudocolored with a custom look-up table spanning from blue via red to yellow hues. Images were exported as RGB images to Photoshop, greyscale images were additionally despeckled.

#### RNA Sequence Data Analysis

Data analysis was completed using Omicsoft Array Studio. On average, each sample was sequenced at a depth of 52 million paired-end reads. Illumina adaptors were stripped during the BCL to FastQ conversion. All raw data was QC’d using the “Raw Data QC” function within Array Studio. Poor quality reads were filtered out using a Q score cutoff of 20. Following the above filter criteria, about 41 million paired-end reads per sample were uniquely mapped to the B38 Mouse Reference Genome using the Ensembl.R90 Gene Model. Raw counts were then converted to FPKM using the “Report Gene/Transcript Counts” function in Array Studio. FPKM was filtered using a cutoff of 0.1 then normalized using the 75th quantile. A constant of 1 was added to all normalized FPKM data prior to a Log2 transformation. This data was used in all downstream analysis including Principal Component Analysis (PCA), heatmaps, TTests and pathway analysis (Ingenuity). Outliers (a total of nine) were confirmed visually using PCA plots as well as Hierarchical Clustering (unsupervised). TTests were generated using the General Linear model in Array Studio. Pathway analysis was completed using fold change and p-values obtained via TTests and imported into Ingenuity (Qiagen).

#### Phagocyte infiltration and activation

Overview images were segmented in cortical layers using a nucleic acid stain (NeuroTrace) and analyzed in VOIs 1000-2000 µm laterally to the midline, ipsi- and contralaterally to the cytokine injection site. In the volumes counted, numbers of CX3CR1^GFP^ cells, co-labelling of CX3CR1^GFP^ and MHCII immunofluorescence and numbers of CCR2^RFP^ cells or Iba1 positive, CX3CR1^GFP^ negative cells, respectively, were analyzed using the cell counter plugin (Fiji). Absolute numbers were adjusted to a standard volume, microglial activation presented as % of MHCII positivity of all CX3CR1^GFP^ cells. For generating the heatmap of inflammation, cells were evaluated in counting frames of 500 x 500 µm size. For accessing the effect of CSF-1R inhibitor compound, three counting frames of 200 x 200 µm per section (six sections per animal) were chosen in layers III-IV at 1000 to 2000 µm lateral to the midline.

#### Analysis of synapse phagocytosis

Ten individual CX3CR1^GFP^ or CCR2^RFP^ cells per animal were reconstructed automatically by Imaris software in a stack of 5.14 µm. The rendered surface was used as a mask on Synapsin-1, PSD-95, Homer-1 staining channel and the spot function was used to identify the positive signals (spheres). Spheres that were not fully engulfed by the phagocytic cell were manually removed. Spheres were checked on LAMP-1 channel to evaluate colocalization. The LAMP1 channel was further masked by the cell surface and the volume was automatically reconstructed to identify the LAMP1 volume per cell.

### Statistical analysis

Sample sizes were chosen according to previous *in vivo* imaging studies (34). Statistical significance was calculated with Prism (Versions 6.0 and 7.0, Graphpad) using ANOVA and t-tests (where normal distribution could be assumed) or Kruskal-Wallis and Mann-Whitney U-test (where non-normal distribution was suspected and confirmed by Shapiro-Wilk test) as described in the figure legends. Data are expressed as the mean ± standard error of the mean. Obtained p-values were corrected for multiple comparisons using Bonferroni’s, Holm-Sidak’sor Dunnett’s procedure and stated as significance levels in the figure legends. In all analyses p<0.05 was considered statistically significant.

### Data and code availability

All data is available in the manuscript or the supplementary materials. Raw data is available upon reasonable request to the corresponding author.

## SUPPLEMENTAL INFORMATION

Figures S1-S8

Movie S1

## REFERENCES

Albert, M., Barrantes-Freer, A., Lohrberg, M., Antel, J.P., Prineas, J.W., Palkovits, M., Wolff, J.R., Brück, W., and Stadelmann, C. (2017). Synaptic pathology in the cerebellar dentate nucleus in chronic multiple sclerosis. Brain Pathol 27, 737–747.

Bialas, A.R., Presumey, J., Das, A., van der Poel, C.E., Lapchak, P.H., Mesin, L., Victora, G., Tsokos, G.C., Mawrin, C., Herbst, R., et al. (2017). Microglia-dependent synapse loss in type I interferon-mediated lupus. Nature 546, 539–543.

Busche, M.A., Eichhoff, G., Adelsberger, H., Abramowski, D., Wiederhold, K.H., Haass, C., Staufenbiel, M., Konnerth, A., and Garaschuk, O. (2008). Clusters of hyperactive neurons near amyloid plaques in a mouse model of Alzheimer’s disease. Science 321, 1686–1689.

Busche, M.A., Wegmann, S., Dujardin, S., Commins, C., Schiantarelli, J., Klickstein, N., Kamath, T.V., Carlson, G.A., Nelken, I., and Hyman, B.T. (2019). Tau impairs neural circuits, dominating amyloid-beta effects, in Alzheimer models in vivo. Nat Neurosci 22, 57–64.

Calabrese, M., Magliozzi, R., Ciccarelli, O., Geurts, J.J.G., Reynolds, R., and Martin, R. (2015). Exploring the origins of grey matter damage in multiple sclerosis. Nat Rev Neurosci 16, 147–158.

Calabrese, M., Poretto, V., Favaretto, A., Alessio, S., Bernardi, V., Romualdi, C., Rinaldi, F., Perini, P., and Gallo, P. (2012). Cortical lesion load associates with progression of disability in multiple sclerosis. Brain 135, 2952–2961.

Chen, T.-W., Wardill, T.J., Sun, Y., Pulver, S.R., Renninger, S.L., Baohan, A., Schreiter, E.R., Kerr, R.A., Orger, M.B., Jayaraman, V., et al. (2013). Ultrasensitive fluorescent proteins for imaging neuronal activity. Nature 499, 295–300.

Confavreux, C., and Vukusic, S. (2006). Age at disability milestones in multiple sclerosis. Brain 129, 595–605.

Damjanovic, D., Valsasina, P., Rocca, M.A., Stromillo, M.L., Gallo, A., Enzinger, C., Hulst, H.E., Rovira, A., Muhlert, N., De Stefano, N., et al. (2017). Hippocampal and Deep Gray Matter Nuclei Atrophy Is Relevant for Explaining Cognitive Impairment in MS: A Multicenter Study. AJNR Am J Neuroradiol 38, 18–24.

Dana, H., Chen, T.W., Hu, A., Shields, B.C., Guo, C., Looger, L.L., Kim, D.S., and Svoboda, K. (2014). Thy1-GCaMP6 transgenic mice for neuronal population imaging in vivo. PLoS One 9, e108697.

DeKosky, S.T., and Scheff, S.W. (1990). Synapse loss in frontal cortex biopsies in Alzheimer’s disease: correlation with cognitive severity. Ann Neurol 27, 457–464.

Di Liberto, G., Pantelyushin, S., Kreutzfeldt, M., Page, N., Musardo, S., Coras, R., Steinbach, K., Vincenti, I., Klimek, B., Lingner, T., et al. (2018). Neurons under T Cell Attack Coordinate Phagocyte-Mediated Synaptic Stripping. Cell.

Dutta, R., Chang, A., Doud, M.K., Kidd, G.J., Ribaudo, M.V., Young, E.A., Fox, R.J., Staugaitis, S.M., and Trapp, B.D. (2011). Demyelination causes synaptic alterations in hippocampi from multiple sclerosis patients. Ann Neurol 69, 445–454.

Ellwardt, E., Pramanik, G., Luchtman, D., Novkovic, T., Jubal, E.R., Vogt, J., Arnoux, I., Vogelaar, C.F., Mandal, S., Schmalz, M., et al. (2018). Maladaptive cortical hyperactivity upon recovery from experimental autoimmune encephalomyelitis. Nat Neurosci 21, 1392–1403.

Elmore, M.R.P., Najafi, A.R., Koike, M.A., Dagher, N.N., Spangenberg, E.E., Rice, R.A., Kitazawa, M., Matusow, B., Nguyen, H., West, B.L., et al. (2014). Colony-stimulating factor 1 receptor signaling is necessary for microglia viability, unmasking a microglia progenitor cell in the adult brain. Neuron 82, 380–397.

Eshaghi, A., Prados, F., Brownlee, W.J., Altmann, D.R., Tur, C., Cardoso, M.J., De Angelis, F., van de Pavert, S.H., Cawley, N., De Stefano, N., et al. (2018). Deep gray matter volume loss drives disability worsening in multiple sclerosis. Ann Neurol 83, 210–222.

Fox, R.J., Coffey, C.S., Conwit, R., Cudkowicz, M.E., Gleason, T., Goodman, A., Klawiter, E.C., Matsuda, K., McGovern, M., Naismith, R.T., et al. (2018). Phase 2 Trial of Ibudilast in Progressive Multiple Sclerosis. N Engl J Med 379, 846–855.

Friese, M.A., Schattling, B., and Fugger, L. (2014). Mechanisms of neurodegeneration and axonal dysfunction in multiple sclerosis. Nat Rev Neurol 10, 225–238.

Gardner, C., Magliozzi, R., Durrenberger, P.F., Howell, O.W., Rundle, J., and Reynolds, R. (2013). Cortical grey matter demyelination can be induced by elevated pro-inflammatory cytokines in the subarachnoid space of MOG-immunized rats. Brain 136, 3596–3608.

Grienberger, C., Rochefort, N.L., Adelsberger, H., Henning, H.A., Hill, D.N., Reichwald, J., Staufenbiel, M., and Konnerth, A. (2012). Staged decline of neuronal function in vivo in an animal model of Alzheimer’s disease. Nat Commun 3, 774.

Hawker, K., O’Connor, P., Freedman, M.S., Calabresi, P.A., Antel, J., Simon, J., Hauser, S., Waubant, E., Vollmer, T., Panitch, H., et al. (2009). Rituximab in patients with primary progressive multiple sclerosis: results of a randomized double-blind placebo-controlled multicenter trial. Ann Neurol 66, 460–471.

Holtmaat, A., Bonhoeffer, T., Chow, D.K., Chuckowree, J., De Paola, V., Hofer, S.B., Hubener, M., Keck, T., Knott, G., Lee, W.C., et al. (2009). Long-term, high-resolution imaging in the mouse neocortex through a chronic cranial window. Nat Protoc 4, 1128–1144.

Hong, S., Beja-Glasser, V.F., Nfonoyim, B.M., Frouin, A., Li, S., Ramakrishnan, S., Merry, K.M., Shi, Q., Rosenthal, A., Barres, B.A., et al. (2016). Complement and microglia mediate early synapse loss in Alzheimer mouse models. Science 352, 712–716.

Hong, S., and Stevens, B. (2016). Microglia: Phagocytosing to Clear, Sculpt, and Eliminate. Dev Cell 38, 126–128.

Huang, D.R., Wang, J., Kivisakk, P., Rollins, B.J., and Ransohoff, R.M. (2001). Absence of monocyte chemoattractant protein 1 in mice leads to decreased local macrophage recruitment and antigen-specific T helper cell type 1 immune response in experimental autoimmune encephalomyelitis. J Exp Med 193, 713–726.

Jürgens, T., Jafari, M., Kreutzfeldt, M., Bahn, E., Brück, W., Kerschensteiner, M., and Merkler, D. (2016). Reconstruction of single cortical projection neurons reveals primary spine loss in multiple sclerosis. Brain 139, 39–46.

Kerschensteiner, M., Bareyre, F.M., Buddeberg, B.S., Merkler, D., Stadelmann, C., Bruck, W., Misgeld, T., and Schwab, M.E. (2004). Remodeling of axonal connections contributes to recovery in an animal model of multiple sclerosis. J Exp Med 200, 1027–1038.

Krasemann, S., Madore, C., Cialic, R., Baufeld, C., Calcagno, N., El Fatimy, R., Beckers, L., O’Loughlin, E., Xu, Y., Fanek, Z., et al. (2017). The TREM2-APOE Pathway Drives the Transcriptional Phenotype of Dysfunctional Microglia in Neurodegenerative Diseases. Immunity 47, 566–581 e569.

Kuchibhotla, K.V., Goldman, S.T., Lattarulo, C.R., Wu, H.-Y., Hyman, B.T., and Bacskai, B.J. (2008). Abeta plaques lead to aberrant regulation of calcium homeostasis in vivo resulting in structural and functional disruption of neuronal networks. Neuron 59, 214–225.

Lagumersindez-Denis, N., Wrzos, C., Mack, M., Winkler, A., van der Meer, F., Reinert, M.C., Hollasch, H., Flach, A., Brühl, H., Cullen, E., et al. (2017). Differential contribution of immune effector mechanisms to cortical demyelination in multiple sclerosis. Acta Neuropathol 134, 15–34.

Lodygin, D., Hermann, M., Schweingruber, N., Flugel-Koch, C., Watanabe, T., Schlosser, C., Merlini, A., Korner, H., Chang, H.F., Fischer, H.J., et al. (2019). beta-Synuclein-reactive T cells induce autoimmune CNS grey matter degeneration. Nature 566, 503–508.

Lublin, F., Miller, D.H., Freedman, M.S., Cree, B.A.C., Wolinsky, J.S., Weiner, H., Lubetzki, C., Hartung, H.P., Montalban, X., Uitdehaag, B.M.J., et al. (2016). Oral fingolimod in primary progressive multiple sclerosis (INFORMS): a phase 3, randomised, double-blind, placebo-controlled trial. Lancet 387, 1075–1084.

Lucchinetti, C.F., Popescu, B.F.G., Bunyan, R.F., Moll, N.M., Roemer, S.F., Lassmann, H., Brück, W., Parisi, J.E., Scheithauer, B.W., Giannini, C., et al. (2011). Inflammatory Cortical Demyelination in Early Multiple Sclerosis. N Engl J Med 365, 2188–2197.

Magliozzi, R., Howell, O.W., Nicholas, R., Cruciani, C., Castellaro, M., Romualdi, C., Rossi, S., Pitteri, M., Benedetti, M.D., Gajofatto, A., et al. (2018). Inflammatory intrathecal profiles and cortical damage in multiple sclerosis. Ann Neurol 83, 739–755.

Mahad, D.H., Trapp, B.D., and Lassmann, H. (2015). Pathological mechanisms in progressive multiple sclerosis. Lancet Neurol 14, 183–193.

Marinkovic, P., Blumenstock, S., Goltstein, P.M., Korzhova, V., Peters, F., Knebl, A., and Herms, J. (2019). In vivo imaging reveals reduced activity of neuronal circuits in a mouse tauopathy model. Brain 142, 1051–1062.

Meijer, K.A., Eijlers, A.J.C., Geurts, J.J.G., and Schoonheim, M.M. (2018). Staging of cortical and deep grey matter functional connectivity changes in multiple sclerosis. J Neurol Neurosurg Psychiatry 89, 205–210.

Menkes-Caspi, N., Yamin, H.G., Kellner, V., Spires-Jones, T.L., Cohen, D., and Stern, E.A. (2015). Pathological tau disrupts ongoing network activity. Neuron 85, 959–966.

Merkler, D., Ernsting, T., Kerschensteiner, M., Brück, W., and Stadelmann, C. (2006). A new focal EAE model of cortical demyelination: multiple sclerosis-like lesions with rapid resolution of inflammation and extensive remyelination. Brain 129, 1972–1983.

Merlini, M., Rafalski, V.A., Rios Coronado, P.E., Gill, T.M., Ellisman, M., Muthukumar, G., Subramanian, K.S., Ryu, J.K., Syme, C.A., Davalos, D., et al. (2019). Fibrinogen Induces Microglia-Mediated Spine Elimination and Cognitive Impairment in an Alzheimer’s Disease Model. Neuron 101, 1099–1108 e1096.

Michailidou, I., Willems, J.G., Kooi, E.J., van Eden, C., Gold, S.M., Geurts, J.J., Baas, F., Huitinga, I., and Ramaglia, V. (2015). Complement C1q-C3-associated synaptic changes in multiple sclerosis hippocampus. Ann Neurol 77, 1007–1026.

Montey, K.L., and Quinlan, E.M. (2011). Recovery from chronic monocular deprivation following reactivation of thalamocortical plasticity by dark exposure. Nat Commun 2, 317.

Nissen, J.C., Thompson, K.K., West, B.L., and Tsirka, S.E. (2018). Csf1R inhibition attenuates experimental autoimmune encephalomyelitis and promotes recovery. Exp Neurol 307, 24–36.

Olmos-Alonso, A., Schetters, S.T., Sri, S., Askew, K., Mancuso, R., Vargas-Caballero, M., Holscher, C., Perry, V.H., and Gomez-Nicola, D. (2016). Pharmacological targeting of CSF1R inhibits microglial proliferation and prevents the progression of Alzheimer’s-like pathology. Brain 139, 891–907.

Ontaneda, D., Thompson, A.J., Fox, R.J., and Cohen, J.A. (2017). Progressive multiple sclerosis: prospects for disease therapy, repair, and restoration of function. Lancet 389, 1357–1366.

Peterson, J.W., Bö, L., Mörk, S., Chang, A., and Trapp, B.D. (2001). Transected neurites, apoptotic neurons, and reduced inflammation in cortical multiple sclerosis lesions. Ann Neurol 50, 389–400.

Reich, D.S., Lucchinetti, C.F., and Calabresi, P.A. (2018). Multiple Sclerosis. N Engl J Med 378, 169–180.

Rocca, M.A., Parisi, L., Pagani, E., Copetti, M., Rodegher, M., Colombo, B., Comi, G., Falini, A., and Filippi, M. (2014). Regional but not global brain damage contributes to fatigue in multiple sclerosis. Radiology 273, 511–520.

Rocca, M.A., Valsasina, P., Absinta, M., Riccitelli, G., Rodegher, M.E., Misci, P., Rossi, P., Falini, A., Comi, G., and Filippi, M. (2010). Default-mode network dysfunction and cognitive impairment in progressive MS. Neurology 74, 1252–1259.

Rodriguez, A., Ehlenberger, D., Kelliher, K., Einstein, M., Henderson, S.C., Morrison, J.H., Hof, P.R., and Wearne, S.L. (2003). Automated reconstruction of three-dimensional neuronal morphology from laser scanning microscopy images. Methods 30, 94–105.

Rose, T., Jaepel, J., Hübener, M., and Bonhoeffer, T. (2016). Cell-specific restoration of stimulus preference after monocular deprivation in the visual cortex. Science 352, 1319–1322.

Saederup, N., Cardona, A.E., Croft, K., Mizutani, M., Cotleur, A.C., Tsou, C.L., Ransohoff, R.M., and Charo, I.F. (2010). Selective chemokine receptor usage by central nervous system myeloid cells in CCR2-red fluorescent protein knock-in mice. PLoS One 5, e13693.

Scalfari, A., Neuhaus, A., Daumer, M., Ebers, G.C., and Muraro, P.A. (2011). Age and disability accumulation in multiple sclerosis. Neurology 77, 1246–1252.

Scalfari, A., Romualdi, C., Nicholas, R.S., Mattoscio, M., Magliozzi, R., Morra, A., Monaco, S., Muraro, P.A., and Calabrese, M. (2018). The cortical damage, early relapses, and onset of the progressive phase in multiple sclerosis. Neurology 90, e2107–e2118.

Schain, A.J., Hill, R.A., and Grutzendler, J. (2014). Label-free in vivo imaging of myelinated axons in health and disease with spectral confocal reflectance microscopy. Nat Med 20, 443–449.

Segal, M., and Korkotian, E. (2014). Endoplasmic reticulum calcium stores in dendritic spines. Front Neuroanat 8, 64.

Siffrin, V., Radbruch, H., Glumm, R., Niesner, R., Paterka, M., Herz, J., Leuenberger, T., Lehmann, S.M., Luenstedt, S., Rinnenthal, J.L., et al. (2010). In vivo imaging of partially reversible th17 cell-induced neuronal dysfunction in the course of encephalomyelitis. Immunity 33, 424–436.

Silva, B.A., Leal, M.C., Farias, M.I., Avalos, J.C., Besada, C.H., Pitossi, F.J., and Ferrari, C.C. (2018). A new focal model resembling features of cortical pathology of the progressive forms of multiple sclerosis: Influence of innate immunity. Brain Behav Immun 69, 515–531.

Siskova, Z., Justus, D., Kaneko, H., Friedrichs, D., Henneberg, N., Beutel, T., Pitsch, J., Schoch, S., Becker, A., von der Kammer, H., et al. (2014). Dendritic structural degeneration is functionally linked to cellular hyperexcitability in a mouse model of Alzheimer’s disease. Neuron 84, 1023–1033.

Tanabe, S., Saitoh, S., Miyajima, H., Itokazu, T., and Yamashita, T. (2019). Microglia suppress the secondary progression of autoimmune encephalomyelitis. Glia 67, 1694–1704.

Terry, R.D., Masliah, E., Salmon, D.P., Butters, N., DeTeresa, R., Hill, R., Hansen, L.A., and Katzman, R. (1991). Physical basis of cognitive alterations in Alzheimer’s disease: synapse loss is the major correlate of cognitive impairment. Ann Neurol 30, 572–580.

Thestrup, T., Litzlbauer, J., Bartholomäus, I., Mues, M., Russo, L., Dana, H., Kovalchuk, Y., Liang, Y., Kalamakis, G., Laukat, Y., et al. (2014). Optimized ratiometric calcium sensors for functional in vivo imaging of neurons and T lymphocytes. Nat Methods 11, 175–182.

Vasek, M.J., Garber, C., Dorsey, D., Durrant, D.M., Bollman, B., Soung, A., Yu, J., Perez-Torres, C., Frouin, A., Wilton, D.K., et al. (2016). A complement-microglial axis drives synapse loss during virus-induced memory impairment. Nature 534, 538–543.

Vogelstein, J.T., Packer, A.M., Machado, T.A., Sippy, T., Babadi, B., Yuste, R., and Paninski, L. (2010). Fast nonnegative deconvolution for spike train inference from population calcium imaging. J Neurophysiol 104, 3691–3704.

Wearne, S.L., Rodriguez, A., Ehlenberger, D.B., Rocher, A.B., Henderson, S.C., and Hof, P.R. (2005). New techniques for imaging, digitization and analysis of three-dimensional neural morphology on multiple scales. Neuroscience 136, 661–680.

Wegner, C., Esiri, M.M., Chance, S.A., Palace, J., and Matthews, P.M. (2006). Neocortical neuronal, synaptic, and glial loss in multiple sclerosis. Neurology 67, 960–967.

Williams, P.R., Marincu, B.-N., Sorbara, C.D., Mahler, C.F., Schumacher, A.-M., Griesbeck, O., Kerschensteiner, M., and Misgeld, T. (2014). A recoverable state of axon injury persists for hours after spinal cord contusion in vivo. Nat Commun 5, 5683.

Witte, M.E., Schumacher, A.M., Mahler, C.F., Bewersdorf, J.P., Lehmitz, J., Scheiter, A., Sanchez, P., Williams, P.R., Griesbeck, O., Naumann, R., et al. (2019). Calcium Influx through Plasma-Membrane Nanoruptures Drives Axon Degeneration in a Model of Multiple Sclerosis. Neuron 101, 615–624 e615.

Wolinsky, J.S., Narayana, P.A., O’Connor, P., Coyle, P.K., Ford, C., Johnson, K., Miller, A., Pardo, L., Kadosh, S., Ladkani, D., et al. (2007). Glatiramer acetate in primary progressive multiple sclerosis: results of a multinational, multicenter, double-blind, placebo-controlled trial. Ann Neurol 61, 14–24.

Wu, Y., Whiteus, C., Xu, C.S., Hayworth, K.J., Weinberg, R.J., Hess, H.F., and De Camilli, P. (2017). Contacts between the endoplasmic reticulum and other membranes in neurons. Proc Natl Acad Sci U S A 114, E4859–E4867.

Xu, T., Yu, X., Perlik, A.J., Tobin, W.F., Zweig, J.A., Tennant, K., Jones, T., and Zuo, Y. (2009). Rapid formation and selective stabilization of synapses for enduring motor memories. Nature 462, 915–919.

Yamasaki, R., Lu, H., Butovsky, O., Ohno, N., Rietsch, A.M., Cialic, R., Wu, P.M., Doykan, C.E., Lin, J., Cotleur, A.C., et al. (2014). Differential roles of microglia and monocytes in the inflamed central nervous system. J Exp Med 211, 1533–1549.

Yang, G., Parkhurst, C.N., Hayes, S., and Gan, W.B. (2013). Peripheral elevation of TNF-alpha leads to early synaptic abnormalities in the mouse somatosensory cortex in experimental autoimmune encephalomyelitis. Proc Natl Acad Sci U S A 110, 10306–10311.

