## Supplemental Materials for "Localized calcium accumulations prime synapses for phagocyte removal in cortical neuroinflammation"

**This PDF file includes:**

fig. S1 to S8
Captions for Movie S1

**Other Supplementary Materials for this manuscript include the following:**

Movie S1

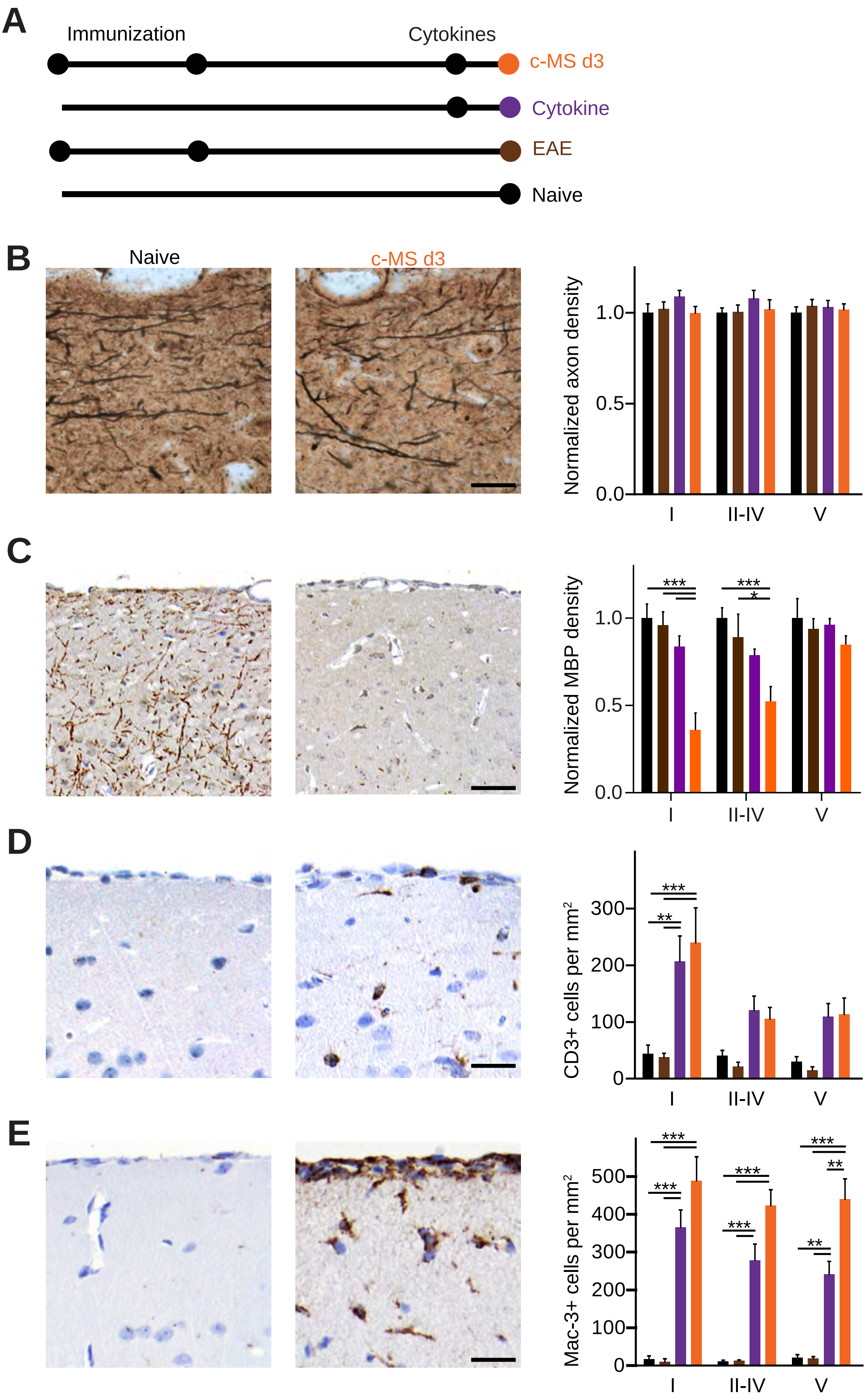

A
Immunization
Cytokines
c-MS d3
Cytokine
EAE
Naive
B
Naive
c-MS d3
Normalized axon density
1.0
0.5
0.0
I
II-IV
V
C
Normalized MBP density
***
*
1.0
0.5
0.0
I
II-IV
V
D
CD3+ cells per mm²
***
**
300
200
100
0
I
II-IV
V
E
Mac-3+ cells per mm²
***
**
500
400
300
200
100
0
I
II-IV
V

**Figure S1 - Histopathological characterization of a mouse model of cortical multiple sclerosis. Related to Figure 1.**

**(A)** Schematic diagram of the experimental design to study histopathological changes in the cortex of age-matched naive (black), EAE (brown), only cytokine injected (violet) and cortical MS model (c-MS d3, orange) mice. **(B)** Bielschowsky's staining was used to analyze axonal density in each layer. Representative images depict Bielschowsky's staining in layer I of naive and c-MS d3 mice (**left, middle,** respectively), axonal density was normalized to average density of the naive group (**right**). **(C)** MBP staining was used to analyze myelin density in each layer. Representative images depict MBP staining in layer I of naive and c-MS d3 mice (**left, middle,** respectively), MBP density was normalized to average density of the naive group (**right**). **(D)** CD3 staining was used to analyze T cells density in each layer. Representative images depict CD3 staining in layer I of naive and c-MS d3 mice (**left, middle,** respectively), CD3 density was normalized to average density of the naive group (**right**). **(E)** Mac-3 staining was used to analyze mononuclear phagocytes density in each layer. Representative images depict Mac-3 staining in layer I of naive and c-MS d3 mice (**left, middle,** respectively), Mac-3 density was normalized to average density of the naive group (**right**). Data shown as mean ± SEM (n= 5, 5, 6 and 6 mice in each group). Scale bars in **B,** 15 µm; **C**, 45 µm **D** and **E,** 25 µm. Two-way RM ANOVA followed by Bonferroni's multiple comparisons test has been performed in **B-E**. *** $P<0.001$, **$P<0.01$, *$P<0.05$.

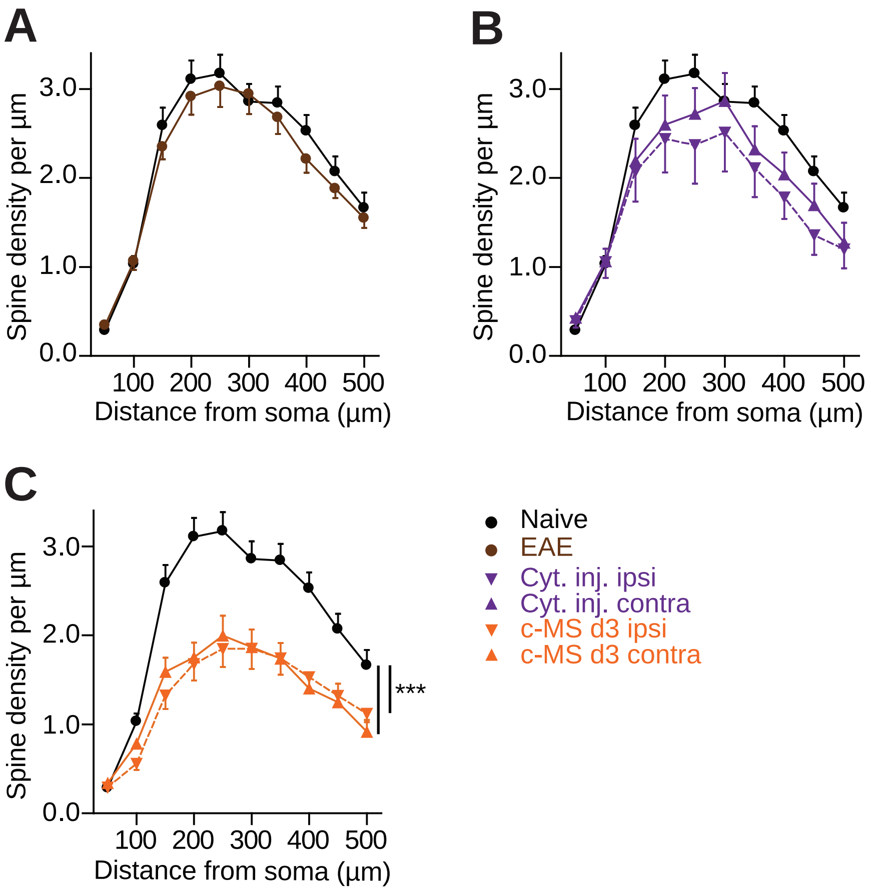
A
B
C
Spine density per µm
Distance from soma (µm)
0.0
1.0
2.0
3.0
100
200
300
400
500
***
Naive
EAE
Cyt. inj. ipsi
Cyt. inj. contra
c-MS d3 ipsi
c-MS d3 contra

**Figure S2 – Induction and spread of spine loss c-MS mice compared to mice with only EAE or sole cytokine injection. Related to Figure 1.**

**(A)** Spine density along the apical dendrite of layer V pyramidal neurons in naive mice and EAE animals without cytokine injection (n=21 neurons from n=7 and 8 mice in each group, respectively; mean ± SEM). **(B)** Spine density along the apical dendrite of layer V pyramidal neurons in naive mice and healthy animals 3d after cytokine injection, ipsi- and contralateral to the injection site (n=21, 8 and 13 neurons from n=7 and 10 mice in each group, respectively; mean ± SEM). **(C)** Spine density along the apical dendrite of layer V pyramidal neurons in naive and c-MS d3 animals, ipsi- and contralateral to the injection site (n=21, 17 and 11 neurons from n=7 and 8 mice in each group, respectively; mean ± SEM). Two-way RM ANOVA followed by Bonferroni's multiple comparisons test has been performed in **A-C**. *** P<0.001.

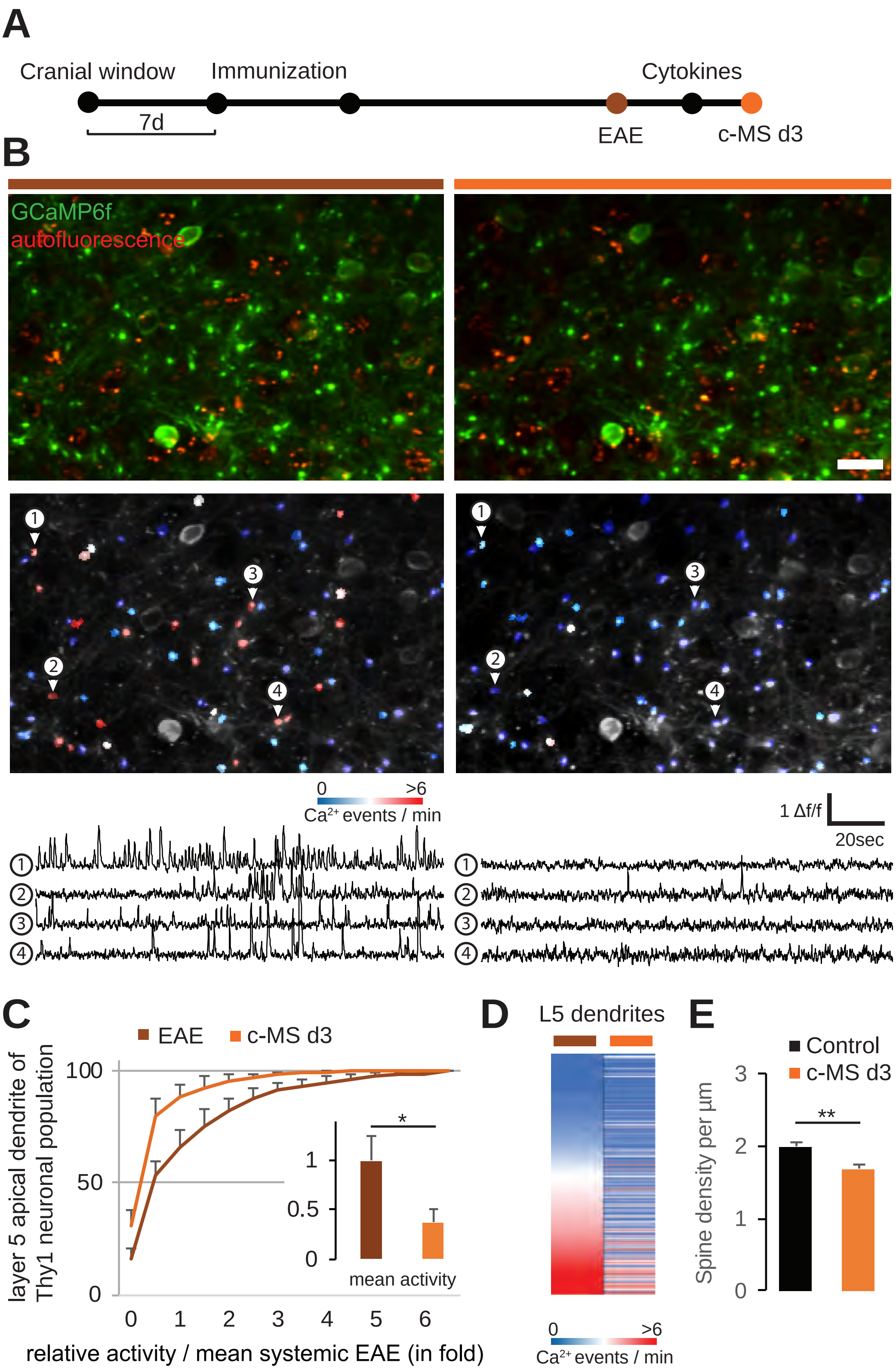
A
Cranial window
Immunization
Cytokines
7d
EAE
c-MS d3
B
GCaMP6f
autofluorescence
0
>6
Ca2+ events / min
1 Δf/f
20sec
C
EAE
c-MS d3
layer 5 apical dendrite of Thy1 neuronal population
100
50
0
1
0.5
0
*
mean activity
0 1 2 3 4 5 6
relative activity / mean systemic EAE (in fold)
D
L5 dendrites
0
>6
Ca2+ events / min
E
Control
c-MS d3
Spine density per µm
3
2
1
0
**

**Figure S3 – Layer 5 projection neurons are silenced in a model of cortical multiple sclerosis. Related to Figure 2.**

**(A)** Schematic diagram of experimental timeline to study longitudinal activity in apical dendrites from somatosensory layer 5 projection neurons in cortical MS model (c-MS), using Thy1-GCaMP6f transgenic animals and *in vivo* imaging. **(B)** *In vivo* multiphoton image of layer 5 apical dendrites over the course of c-MS through a cranial window; **left,** before cytokine injection (EAE) and **right** 3 days (c-MS d3) post cytokine injection. **Top**: Merged GCaMP6f and autofluorescence channels allowing the identification of apical dendrites (green only). **Middle**: GCaMP6f channel masked and color-coded for cytoplasmic $Ca^{2+}$ events per minute. **Bottom**: Representative calcium activity traces displayed as delta f/f for the four apical dendrites marked in the middle panel. **(C)** Cumulative plots representing activity for the entire population of apical dendrites for each selected time point over the course of the cortical MS model. **Inserts**: Mean activity normalized to baseline for EAE and c-MS d3 shown as mean ± SEM (n=3 c-MS mice, Shapiro-Wilk and paired t-test). **(D)** Heat map representation of ranked activities at a single dendrite resolution between EAE and c-MS d3. **(E)** Spine density of dendritic stretches of pyramidal layer V neurons from c-MS animals used in B-D (imaged area was used for quantification) and in control animals (n=30 and n=40 dendritic stretches from n=3 and n=4 mice, respectively; mean ± SEM, unpaired t-test). Scale bar in **B**, 20µm. **P < 0.01, *P<0.05.

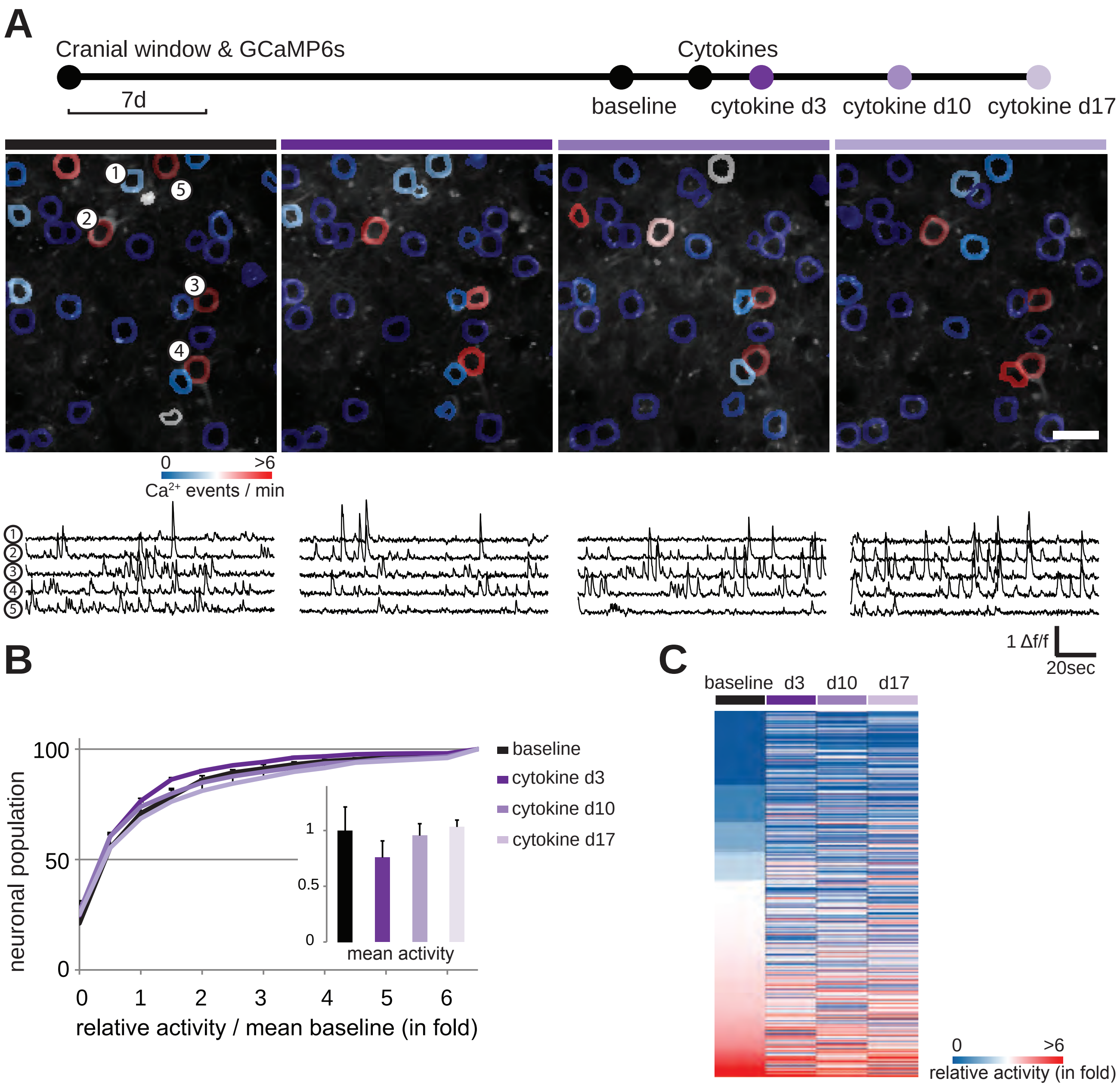
A
Cranial window & GCaMP6s
Cytokines
7d
baseline
cytokine d3
cytokine d10
cytokine d17
0
>6
Ca2+ events / min
1 Δf/f
20sec
B
neuronal population
100
50
0
0 1 2 3 4 5 6
relative activity / mean baseline (in fold)
1
0.5
0
mean activity
baseline
cytokine d3
cytokine d10
cytokine d17
C
baseline d3 d10 d17
0
>6
relative activity (in fold)

**Figure S4 - Recording the activity of single cortical neurons over time in the intact and inflamed gray matter. Related to Figure 2.**

**(A) Top:** Schematic diagram showing longitudinal *in vivo* imaging of $Ca^{2+}$ transients in layer 2/3 somatosensory neurons upon injection of cytokines in healthy animals, using GCaMP6s encoded calcium indicator delivered via viral vector (AAV1.hSyn1.mRuby2.GCaMP6s). **Middle, Bottom:** Longitudinal *in vivo* multiphoton images of the same layer 2/3 neuronal population before and after cytokine injection; from **left** to **right** respectively: baseline, 3 days, 10 days and 17 days post cytokine injection. **Middle**: Grayscale images of GCaMP6s channel masked and color-coded for cytoplasmic $Ca^{2+}$ events per minute. **Bottom**: Representative $Ca^{2+}$ activity traces displayed as delta f/f for the neurons marked in the left panel. **(B)** Cumulative plots of the neuronal activity for the entire population of neurons investigated over the course of the cytokine injection in healthy animals. **Insert**: Mean neuronal activity normalized to baseline for each time point shown as mean ± SEM (n=4 cytokine injected mice, tested with Shapiro-Wilk and one-way ANOVA). **(C)** Heat map representation of the single neurons' activities before and after cytokine injection. Scale bar in **A**, 20μm.

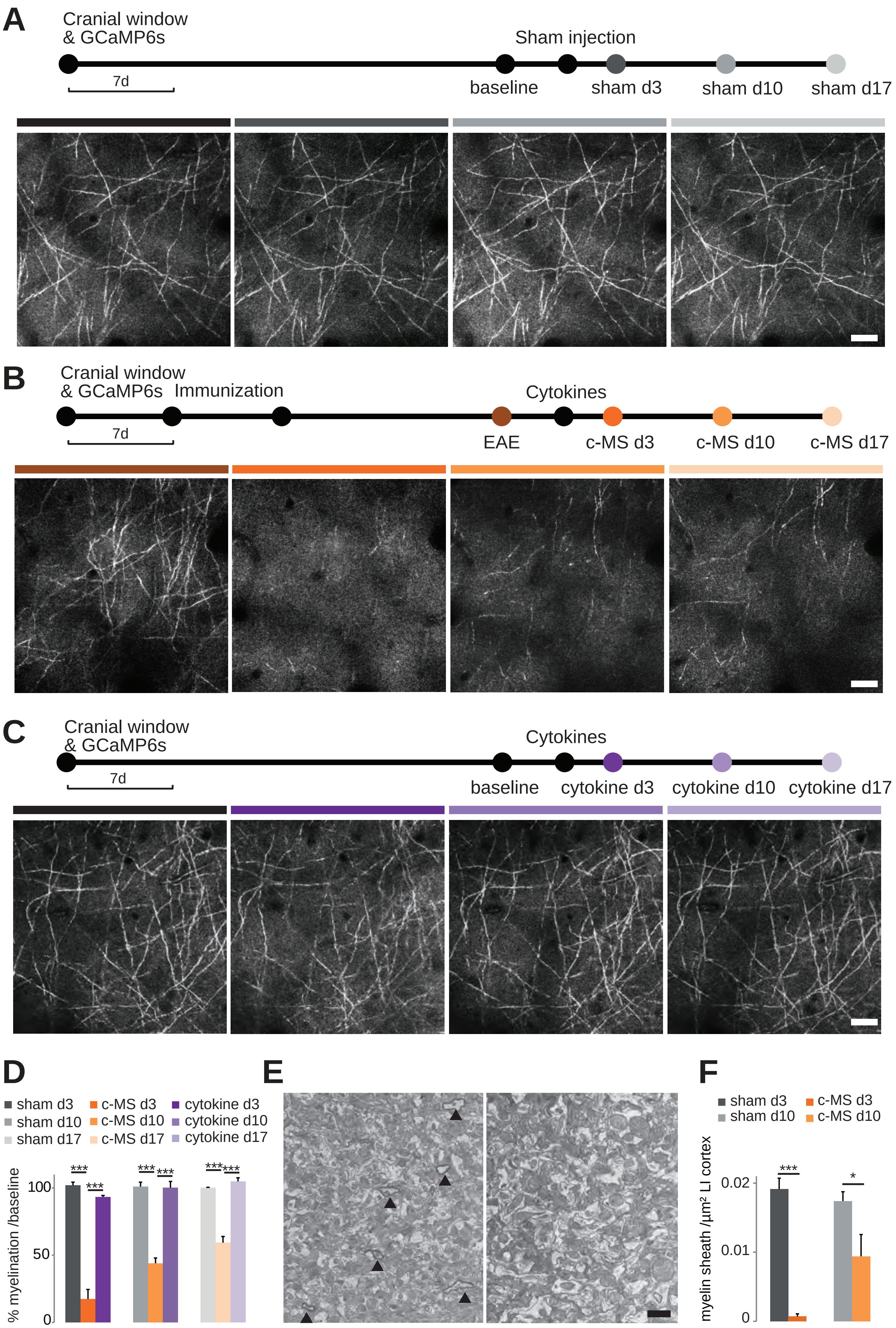
A
Cranial window & GCaMP6s
Sham injection
7d
baseline
sham d3
sham d10
sham d17
B
Cranial window & GCaMP6s
Immunization
Cytokines
7d
EAE
c-MS d3
c-MS d10
c-MS d17
C
Cranial window & GCaMP6s
Cytokines
7d
baseline
cytokine d3
cytokine d10
cytokine d17
D
sham d3
sham d10
sham d17
c-MS d3
c-MS d10
c-MS d17
cytokine d3
cytokine d10
cytokine d17
% myelination /baseline
100
50
0
***
***
E
F
sham d3
sham d10
c-MS d3
c-MS d10
myelin sheath /µm² LI cortex
0.02
0.01
0
***
*

**Figure S5 – Myelination recovers slower than neuronal activity. Related to Figure 2.**

**(A-C) Top**: Schematic diagram showing longitudinal *in vivo* SCoRe imaging of myelin within the top 50 µm of layer 1 somatosensory cortex in sham injected animals **(A)**, animals with cortical MS model **(B)** and in cytokine injected healthy animals **(C)**. **(A-C) Bottom**: Longitudinal *in vivo* SCoRe images of the same layer 1 area. From **left** to **right** respectively: **(A)** baseline, 3 days, 10 days and 17 days post sham injection. **(B)** EAE baseline, 3 days, 10 days and 17 days post cytokine injection. **(C)** baseline, 3 days, 10 days and 17 days post cytokines injection. **(D)** Percentage of SCoRe signal compared to baselines for sham, cytokine only injection and systemic EAE for the c-MS model, 3 days, 10 days and 17 days post injection shown as mean ± SEM (n=3 for sham, cytokine only and c-MS mice, Shapiro-Wilk and One-way ANOVA followed by Bonferroni's multiple comparisons test has been performed). **(E)** Electron micrographs of sham injected (**left**) and c-MS model (**right**) cortex layer 1 showing myelin sheath (black arrowheads) 3 days post injection. **(F)** Myelin sheath density in layer 1 of somatosensory cortex in sham injected and c-MS animals at 3 days and 10 days post injection shown as mean ± SEM (tested in n=4 sham injected and n=4 c-MS mice for day 3 and n=5 sham injected and n=3 c-MS mice for day 10, Shapiro-Wilk and One-way ANOVA followed by Bonferroni's multiple comparisons test). Scale bar in **A**, **B, C**, 20µm; **E**, 1µm.***P<0.001, *P<0.05

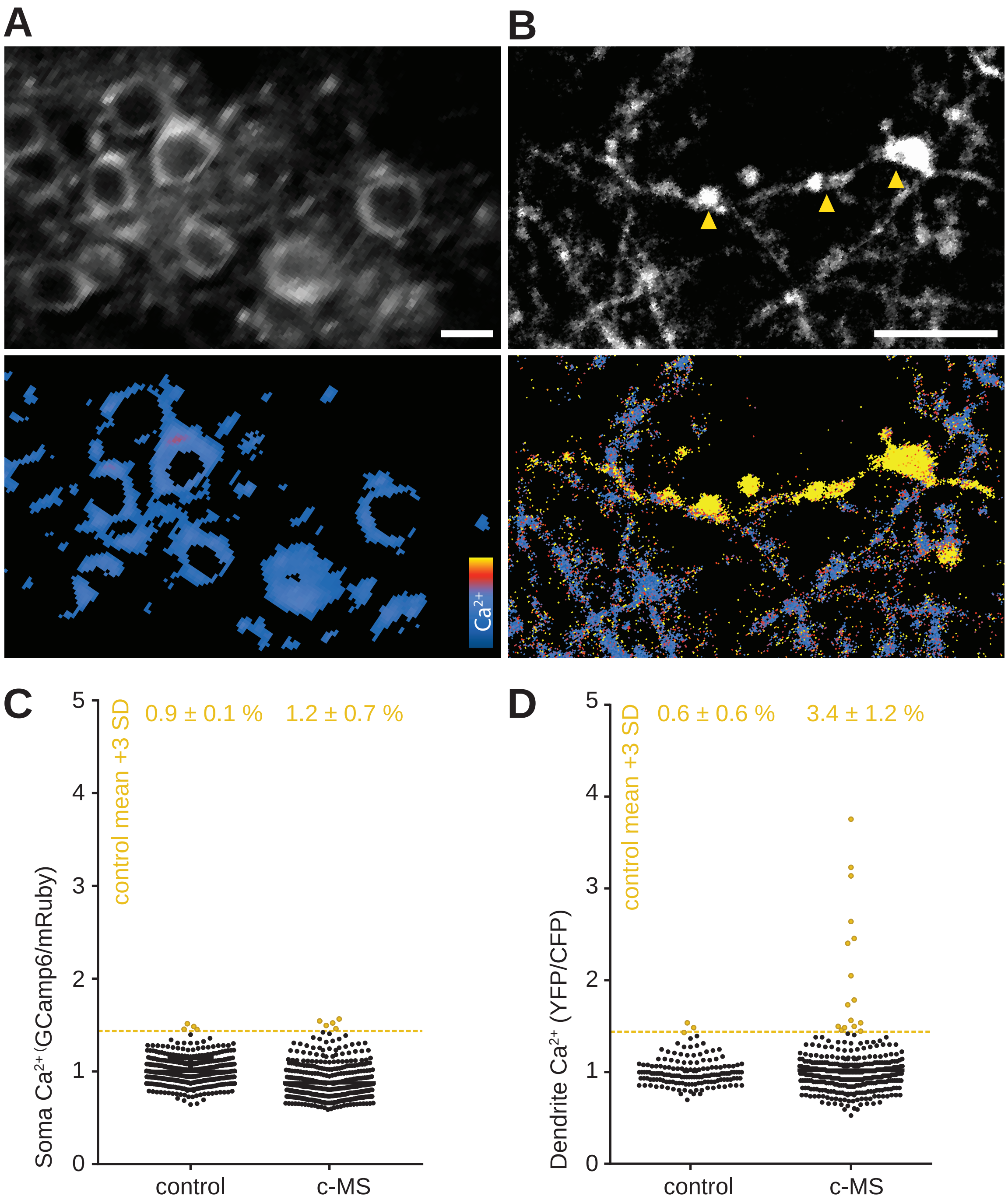
A
B
Ca2+
C
D
5
4
3
2
1
0
control mean +3 SD
0.9 ± 0.1 %
1.2 ± 0.7 %
Soma Ca2+ (GCamp6/mRuby)
control
c-MS
0.6 ± 0.6 %
3.4 ± 1.2 %
Dendrite Ca2+ (YFP/CFP)

**Figure S6 – Ratiometric imaging of calcium levels in the somata and dendrites of cortical neurons. Related to Figure 4.**

**(A)** *In vivo* multiphoton projection images of layer 2-3 neuronal somata and their $Ca^{2+}$ levels in acute cortical MS model. Neurons were labelled using the viral vector AAV1.hSyn.mRuby2.Gcamp6s (see Methods), allowing for ratiometric readout (GCamp6s/mRuby) of baseline neuronal $Ca^{2+}$ concentration. Grayscale images of GCamp6s channel (**top**), ratiometric (GCamp6s/mRuby) images masked and color-coded for cytoplasmic $Ca^{2+}$ (**bottom**). **(B)** *In vivo* multiphoton projection images of apical tuft dendrites and their $Ca^{2+}$ levels in acute c-MS model, labelled with AAV1.hSyn1.Twitch2b. Grayscale images of YFP channel (**top**), ratiometric (YFP/CFP) images masked and color-coded for cytoplasmic $Ca^{2+}$ (**bottom**). Example of a high $Ca^{2+}$ dendrite exhibiting swellings (**arrow heads**). **(C)** $Ca^{2+}$ concentration of single somata in control (sham injection) and c-MS (d3) animals, plotted as GCamp6s/mRuby channel ratios normalized to control mean. **Top:** Percentage of somata per animal with $Ca^{2+}$ concentration > 3SD above control mean, shown as mean ± SEM (tested per animal in n=2 control and n=3 c-MS mice, Mann-Whitney U test). **(D)** $Ca^{2+}$ concentration of single dendrites in healthy control and c-MS d3 animals, plotted as YFP/CFP channel ratios normalized to control mean. **Top:** Percentage of dendrites per animal with $Ca^{2+}$ concentration >3SD above control mean, shown as mean ± SEM (tested per animal in n=8 control and n=17 c-MS mice, Mann-Whitney U test). Scale bars in **A, B**, 10 μm.

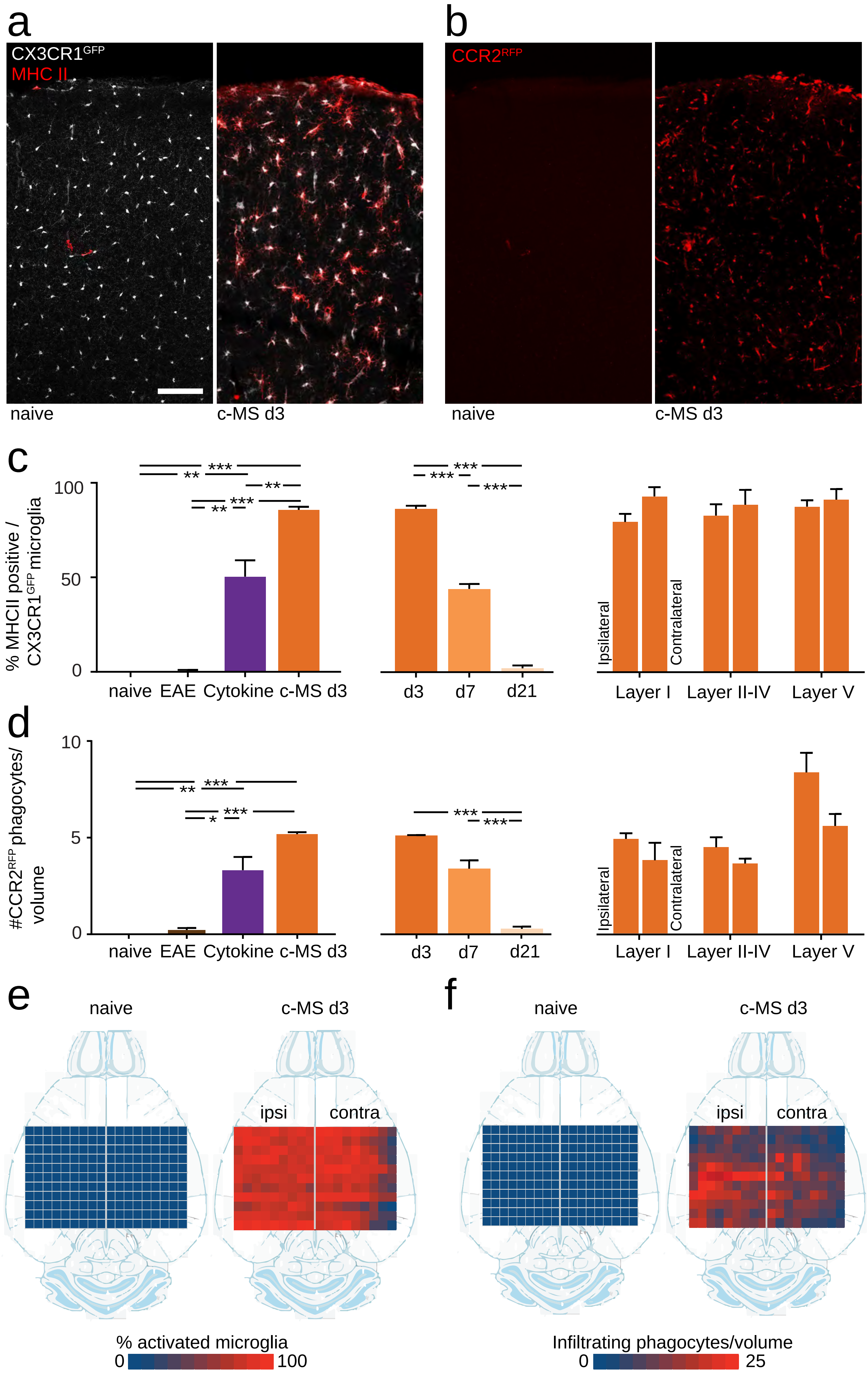
a
CX3CR1GFP
MHC II
naive
c-MS d3
b
CCR2RFP
naive
c-MS d3
c
% MHCII positive / CX3CR1GFP microglia
100
50
0
naive
EAE
Cytokine
c-MS d3
d3
d7
d21
Ipsilateral
Contralateral
Layer I
Layer II-IV
Layer V
d
#CCR2RFP phagocytes/ volume
10
5
0
naive
EAE
Cytokine
c-MS d3
d3
d7
d21
Ipsilateral
Contralateral
Layer I
Layer II-IV
Layer V
e
naive
c-MS d3
ipsi
contra
% activated microglia
0
100
f
naive
c-MS d3
ipsi
contra
Infiltrating phagocytes/volume
0
25

**Figure S7 – Temporal and spatial characteristics of monocyte infiltration and microglial activation in the c-MS model. Related to Figure 5.**

**(A, B)** Confocal projection images of a cortical column in healthy control (naive, **left**) and acute cortical MS model (c-MS d3, **right**) *BiozziABH CX3CR1$^{GFP}$ x CCR2$^{RFP}$* mice. **(A)** Immuno-staining for MHCII (red), as a marker of microglial activation, overlayed on the GFP signal of resident microglia (greyscale). **(B)** RFP signal of blood-born infiltrating phagocytes shown in red. **(C) Left:** Percentages of activated microglia (% MHCII positive/ all CX3CR1$^{GFP}$ cells) in healthy control (naive), only immunized (EAE), only cytokine injected (cytokine) and acute cortical MS model (c-MS d3) mice. **Middle:** Timecourse of microglial activation (in %) 3d, 7d and 21d after induction of cortical MS model (d3, d7, d21). **Right:** Distribution of microglial activation (in %) within the layers of the cortical column (Layer1, Layer 2-4 and Layer 5) and between injected (ipsilateral) and non-injected hemisphere (contralateral). c-MS d3 data plotted in all 3 panels. Shown as mean ± SEM (tested per animal in n=2 naive, n=2 EAE, n=3 Cytokine, n=3 c-MS d3; 1way ANOVA). **(D) Left:** Numbers of infiltrating phagocytes (CCR2$^{RFP}$ cells) per standardized volume in healthy control (naive), only immunized (EAE), only cytokine injected (cytokine) and acute cortical MS model (c-MS d3) mice. **Middle:** Timecourse of phagocyte infiltration 3d, 7d and 21d after induction of cortical MS model (d3, d7, d21). **Right:** Distribution of phagocyte infiltration within the layers of the cortical column (Layer1, Layer 2-4 and Layer 5) and between injected (ipsilateral) and non-injected hemisphere (contralateral). c-MS d3 data plotted in all 3 panels. Shown as mean ± SEM (tested per animal in n=2 naive, n=2 EAE, n=3 Cytokine, n=3 c-MS d3; 1way ANOVA). **(E)** Color-coded spatial matrix representing microglial activation (in %) of healthy control (naive, n=3, **left**) and acute cortical MS model (c-MS d3, n=3, **right**) mice, quantified in 11x8 cortical segments ipsi- and contralateral to the cytokine injection site. **(F)** Color-coded matrix showing numbers infiltrating phagocytes (Iba1 immunolabelling positive, CX3CR1$^{GFP}$ negative cells) in healthy control (naive, n=3, **left**) and acute cortical MS model (c-MS d3, n=3, **right**) mice per cortical segment ipsi- and contralateral to injection site. Scale bar in **A** (applies to B) 100 µm. ***$P<0.001$, **$P<0.01$, *$P<0.05$.

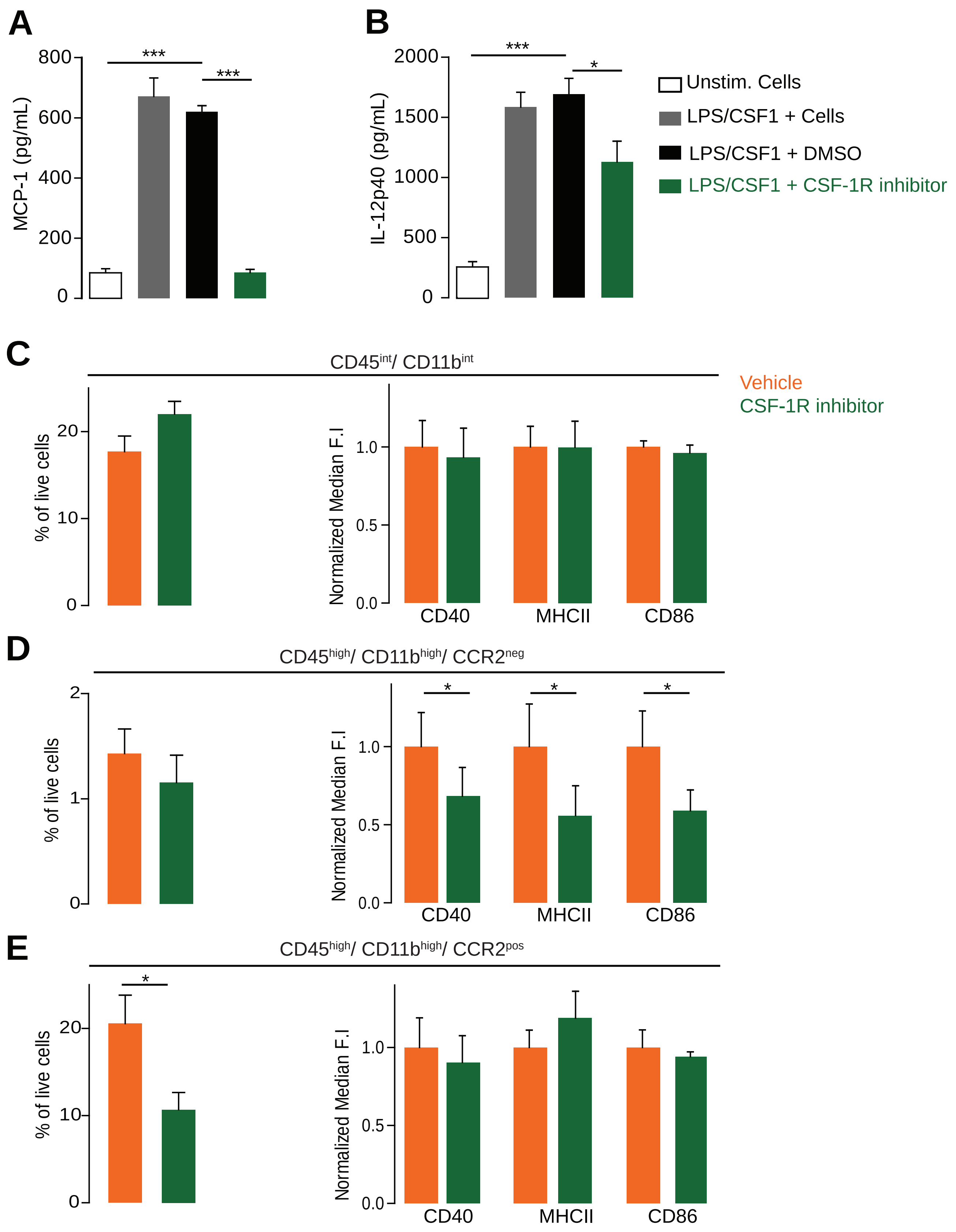
A
MCP-1 (pg/mL)
800
600
400
200
0
***
***
B
IL-12p40 (pg/mL)
2000
1500
1000
500
0
***
*
Unstim. Cells
LPS/CSF1 + Cells
LPS/CSF1 + DMSO
LPS/CSF1 + CSF-1R inhibitor
C
CD45int/ CD11bint
Vehicle
CSF-1R inhibitor
% of live cells
20
10
0
Normalized Median F.I
1.0
0.5
0.0
CD40
MHCII
CD86
D
CD45high/ CD11bhigh/ CCR2neg
% of live cells
2
1
0
Normalized Median F.I
1.0
0.5
0.0
*
*
*
CD40
MHCII
CD86
E
CD45high/ CD11bhigh/ CCR2pos
*
% of live cells
20
10
0
Normalized Median F.I
1.0
0.5
0.0
CD40
MHCII
CD86

**Figure S8 – *In vitro* and *in vivo* effects of CSF-1R inhibitor on inflammation. Related to Figure 6.**

**(A)** Isolated primary murine microglia (untreated) show elevated MCP-1 (CCL2) after treatment with CSF-1. While adding DMSO does not change the levels of MCP-1, CSF-1R inhibitor compound decreases the levels of MCP-1 in primary murine cells (n=3 in all groups, one-way ANOVA). **(B)** Isolated primary murine microglia (untreated) show elevated Il-12p40 after treatment with LPS. While adding DMSO does not change the levels of Il-12p40, CSF-1R inhibitor decreases the levels of Il-12p40 in primary murine cells (n=6 for untreated cells and n=5 for the other groups, one-way ANOVA). (**C-E**) FACS analysis of cell populations in brains of acute c-MS model mice. **(C)** The proportion of CD45 int/ CD11b int cells to all the live cells remained unchanged following CSF-1R inhibitor treatment in the c-MS model compared to vehicle-treated c-MS model (**left**). CD40, MHCII and CD86 markers showed unchanged levels of median fluorescence intensity (**right**). (**D**) The proportion of CD45 high/ CD11b high/ CCR2$^{RFP}$ negative cells to all the live cells remained unchanged following CSF-1R inhibitor treatment in the c-MS model compared to vehicle-treated c-MS model (**left**). CD40, MHCII and CD86 markers revealed differences between the levels of median fluorescence intensity of the two groups (**right**). (**E**) The proportion of CD45 high/ CD11b high/ CCR2 positive cells to all the live cells was decreased following CSF-1R inhibitor treatment in the c-MS model compared to vehicle-treated c-MS model (**left**). CD40, MHCII and CD86 markers showed unchanged levels of median fluorescence intensity (**right**) (n=10 mice were analyzed in each group. Unpaired two-tailed t-test and two-way ANOVA in left panels and right panels of **C-E,** respectively). *** P<0.001, *P<0.05.

**Captions for Movie S1 - Activity of single cortical neurons over the course of cortical MS model.**

Movie generated using FIJI software from 15 Hz recordings and displayed at 100 frames per second with a walking average of 3 (60 second recordings per time point).
